# RNA Quality Control Enables Antibiotic Tolerance

**DOI:** 10.64898/2026.02.02.703294

**Authors:** Andrew T. Nishimoto, Juan C. Ortiz-Marquez, Michelle R. Scribner, Yanying Yu, Qidong Jia, Haley Echlin, Amy R. Iverson, Abigail E. McKnight, Nadia Olivero, Aaron Poole, Enolia Marr, Jordan Coggins, Adam Rosenthal, Chrispin Chaguza, Randolph K. Larsen, Mark E. Hatley, Ralph R. Isberg, Vaughn S. Cooper, Tim van Opijnen, Jason W. Rosch

## Abstract

The mechanisms underlying adaptation to antibiotic pressure within complex host environments remain incompletely understood. By experimentally evolving *Streptococcus pneumoniae* subjected to various antibiotics and immune states, we demonstrate populations adopting distinct adaptive strategies depending on specific selective context. General antibiotic stress drives convergent mutations in *rny*, encoding the RNA degradosome scaffold RNase Y, that exhibit broad-spectrum antibiotic tolerance and accelerated recovery. Single-cell transcriptomics revealed antibiotic-induced death is driven by transcriptional collapse, a catastrophic loss of RNA quantity and integrity. In contrast, *rny* mutants avert this via a bet-hedging strategy: a resilient minority maintains a baseline transcriptional profile, while a quiescent majority undergoes selective RNA degradation to preserve transcript fidelity. Upon stress removal, these populations execute a prioritized transcriptional ribosomal reboot, facilitating accelerated recovery. These findings identify RNA turnover as a tunable master regulator to survive combined pressures of antibiotics and immunity.

## INTRODUCTION

Successful antimicrobial therapy relies on a collaboration between the drug and the host immune system that constrains and ultimately eliminates the pathogen. While the immune system is critical for clearing infection, its role in shaping the evolutionary trajectory of the pathogen, particularly in the emergence of antimicrobial resistance (AMR) and tolerance, remains a black box in infection biology. Standard models of resistance evolution^1,2^ do not account for effects of immune surveillance and in particular how the magnitude and quality of immune pressure might constrain or accelerate the acquisition of pathogen survival mechanisms. This gap is problematic given that immunodeficiency is a critical risk factor for infection^3^. Recent work suggests that the specific immune deficiencies, such as neutropenia, can broaden the landscape of accessible escape mutations in *Acinetobacter baumannii*^4^, whereas functional immunity may constrain evolutionary trajectories in *Pseudomonas aeruginosa* by limiting population density.^5^ Nonetheless, greater understanding of how antimicrobials and host immunity interact to either censor or facilitate the genomic and phenotypic adaptations that lead to treatment failure is needed.

Classically, AMR is defined by stable genetic changes that elevate the minimum inhibitory concentration (MIC). However, these mutations frequently impose fitness costs that slow pathogen growth and may render them more vulnerable to immune clearance^6-9^. Considerable evidence indicates that treatment failure arises not from frank resistance but rather from antibiotic tolerance^10-12^, a phenomenon characterized by prolonged transient survival above the MIC^13^. While tolerance is increasingly recognized as a precursor to resistance^14-18^, its emergence has rarely been systematically evaluated across disparate host immune states. Host factors and the immune response during infection are known to influence bacterial populations, for example by increasing phenotypic heterogeneity and inducing persister cell formation in *Mycobacterium tuberculosis* and *Salmonella typhimurium*, respectively ^19,20^. These findings raise the hypothesis that the host immune status serves as a critical determinant of pathways to pathogen adaptation, selecting for specific modes of resilience that allow survival without the prohibitive costs of classical resistance.

To test this hypothesis, we experimentally evolved *Streptococcus pneumoniae* in the presence of each of 10 antibiotics in mice with three distinct immune states. We systematically mapped their evolutionary dynamics, measured their phenotypic responses to antibiotics, and identified mechanisms causing antibiotic treatment failure. Remarkably, evolution in this antibiotic-treated murine model rarely selected lineages with canonical resistance. Instead, distinct strategies emerged: neutrophil-replete environments selected for specific immune-evasion mutations (e.g., in the nicotinamidase SP_1583), while general antibiotic pressure drove convergent mutations in *rny*, encoding the RNA degradosome scaffold RNase Y whose product coordinate transcript degradation. Specific nonsynonymous *rny* SNPs confer a broad stress tolerance phenotype that is characterized by minimized lag time following drug exposure and rapid population recovery. These discoveries motivated a detailed study of how heterogeneity in the transcriptional program evolves in *S. pneumoniae* mutants arising under antibiotic treatment *in vivo*.

By integrating temporal single-cell RNA-Seq with global RNA integrity analysis, we delineate the molecular mechanism underlying this adaptive, tolerance state. We show that antibiotic-induced cell death in *S. pneumoniae* is driven by transcriptional collapse, a chaotic, high-entropy state characterized by uncontrolled transcription followed by a catastrophic loss of RNA quantity and integrity. In contrast, the evolved *rny* mutants avert this fate via a dual survival strategy: most cells enter a quiescent, high-fidelity RNA state, while a resilient minority maintains a homeostatic baseline, poised for immediate reactivation. These data suggest that lethal stress triggers a systemic failure of transcriptional order, and that pathogens can avoid this fate through a general tolerance response accessed by modulating RNA turnover.

## RESULTS

### Experimental evolution of *S. pneumoniae* Under Diverse Antibiotic Regimens *in vivo*

To evaluate how antibiotic treatment and host immune condition interact to constrain or drive pathogen adaptation, we propagated *S. pneumoniae* populations in mice across three distinct immune states: immunocompetent, neutrophil-depleted, and macrophage-depleted (**Figure 1A**). These lineages were subjected to treatment with one of ten antibiotics spanning multiple classes, administered under three distinct regimens: constant subinhibitory pressure (azithromycin, AZM; ciprofloxacin, CIP; imipenem, IPM), constant-then-escalating pressure (cefepime, CEF; levofloxacin, LVX; linezolid, LNZ), or strictly escalating pressure (meropenem, MEM; penicillin, PEM; rifampicin, RIF; vancomycin VNC) (**Figure S1A, S1B**). This represents data from >5,000 bacterial generations from populations evolved during active murine infection under dual antibiotic and immune selection. For all antibiotics, the initial dose was determined empirically by the antibiotic dose that reduced pneumococcal titers by 90% compared to untreated controls (**Figure S1A**). We monitored bacterial population sizes at each passage by counting viable CFU. As predicted, and consistent with prior observations with *Acinetobacter baumannii* propagated in ciprofloxacin-treated mice^4^, lineages propagated in immune-depleted hosts maintained higher airway population sizes than those in immunocompetent hosts. However, in contrast to this study, whereas neutrophil-depleted *A. baumannii* populations eventually evolved resistance that enabled populations to expand up to 1,000-fold^4^, *S. pneumoniae* populations did not exhibit comparable expansion, even in immunodeficient backgrounds (**Figure S1B**). Critically, despite increasing antibiotic dosages in seven of the ten regimens, bacterial populations were not eradicated but maintained consistent titers following challenge and treatment. These observations suggested that the bacteria were using a survival strategy distinct from classical resistance.

**Figure 1.**
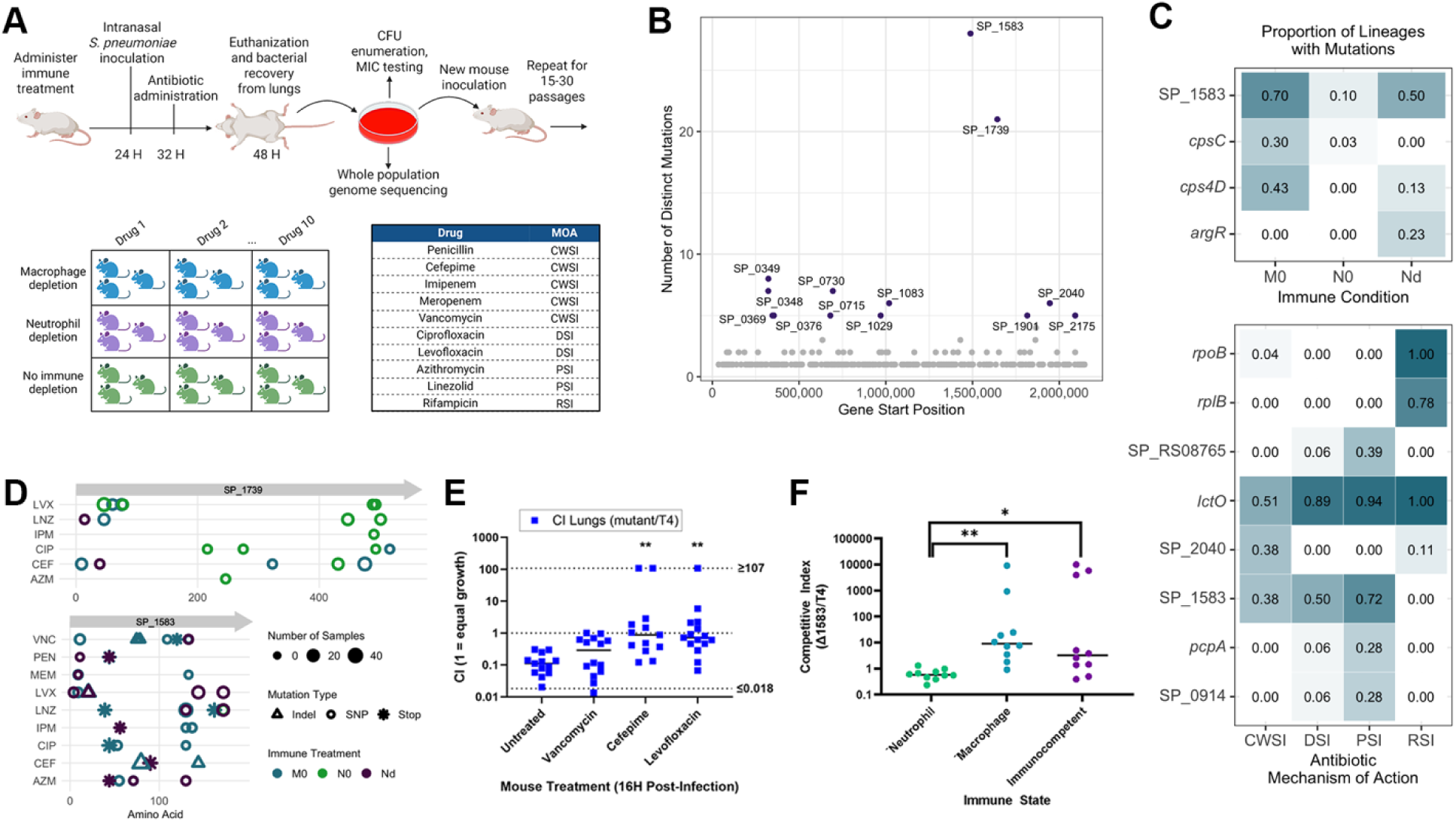
Adaptations to antibiotics and immune status during experimental evolution. (A) Outline of the experimental evolution workflow, the immune depletions being investigated, and the antibiotics used. CWSI- cell wall synthesis inhibitor, PSI- protein synthesis inhibitor, RSI, RNA synthesis inhibitor PSI = protein synthesis inhibitor, CWSI = cell wall synthesis inhibitor, DSI = DNA synthesis inhibitor, RSI = RNA synthesis inhibitor. M0 = macrophage-depleted mouse lineage, N0 = neutrophil-depleted mouse lineage, Nd = non-depleted (wildtype) mouse lineage, VNC = vancomycin, PEN = penicillin, MEM = meropenem, LNZ = linezolid, LVX = levofloxacin, IPM = imipenem, CIP = ciprofloxacin, CEF = cefepime, AZM = azithromycin. (B) The number of independent mutations detected in every gene within the *S. pneumoniae* genome is shown at the start position of the gene within the *S. pneumoniae* genome. Genes that acquired 5 or more mutations are indicated in black and labeled. (C) Proportion of lineages with frequently mutated genes in response to both host immune status and antibiotic mechanism of action. (D) Positions of mutations within the two genes which acquired the most independent mutations, SP_1739 and SP_1583. Point shapes indicate the immune status of the mouse lineage, point size indicates the number of samples within the drug and immune treatment combination that possessed the mutation, and the point shape indicates the mutation type. M0 = macrophage-depleted mouse lineage, N0 = neutrophil-depleted mouse lineage, Nd = non-depleted (wildtype) mouse lineage. (E) The competitive index between the RNAse Y deletion mutant and the parent TIGR4 strain in the lungs of antibiotic-treated or untreated mice is shown. (F) Competitive index the SP_1583 mutant in non-depleted and either neutrophil or mancrophage dpleted mice. Competitive index was calculated as the CFU counts of the mutant divided by the CFU of TIGR4. A competitive index of less than one represents an advantage for the wildtype TIGR4 strain. *= p<0.05, ** = p<0.01 by Mann-Whitney.

### Evolutionary dynamics show interactions between antibiotic-associated mutations and immune status

Immunodeficient mice supported larger *S. pneumoniae* population sizes under all drug treatments (**Figure S1B**), which enables lineages to sample more mutations during their evolution. A central question is whether these lineages consequently evolved more numerous or different adaptations to these conditions. We performed whole-genome population sequencing at every passage to a depth sufficient to identify all mutations reaching 10% frequency or greater. We applied a rigorous filtering strategy to identify high-confidence mutations that reached these frequencies due to direct or linked selection, retaining only those reaching cumulative frequencies across samples of at least 100% per 15-passage span or 200% per 30-passage span (**Figure S1C, S1D**). Lineages accumulated an average of 15 mutations that exceeded these thresholds but were remarkably diverse in total mutation count (minimum = 3, maximum = 47) and gene identities. Populations evolved in neutrophil-depleted lines accumulated the most mutations (491), followed by macrophage-depleted lines (470), and those evolved in wild-type mice accumulated the fewest (404). Genetic similarity was greatest among replicates within a given drug-by-immune-status treatment but was surprisingly low at the level of host immune state or antibiotic class (**Figure S2**). Notable exceptions included lineages evolved in rifampicin, each of which acquired resistance mutations in the drug target *rpoB*. However, for all other antibiotic classes, the genomic landscape was largely devoid of fixed on-target resistance mutations. In most cases, mutations in expected targets, such as penicillin-binding proteins that would confer resistance to β-lactam drugs, failed to sweep the population and were ultimately outcompeted. Nonetheless, mutations were distributed non-randomly and 13 genes acquired five or more independent mutations across lineages, far exceeding the probability of chance accumulation (**Figure S1E, S1F, S3**). In summary, strong selection favored numerous mutations predicted to balance host fitness with antibiotic and immune pressure, independent of classical resistance mechanisms.

### Genetic adaptations to antibiotic-by-immunity interactions

We next evaluated whether mutations in certain genes were significantly more frequent under specific immune or antibiotic pressures (**Figure 1B, 1C**). Four genes were strongly associated with mouse immune status: SP_1583 (nicotinamidase), *argR*, and the capsule biosynthesis genes *cpsC* and *cps4D*. Capsule mutations that are expected to reduce capsule production were notably enriched in macrophage-depleted lineages, consistent with the capsule’s primary role in evading phagocytosis^22-24^. Conversely, mutations in *argR*, a regulator of arginine biosynthesis and virulence gene expression^25,26^, appeared exclusively in immunocompetent hosts. After grouping treatments by antibiotic mechanism of action (inhibition of (i) cell wall synthesis, (ii) DNA synthesis, (iii) RNA synthesis, and (iv) protein synthesis), we identified eight mutated genes (SP_1961 [*rpoB*], SP_0212 [*rplB*], SP_0715 [*lctO*], SP_1769, SP_2040, SP_1583, SP_2136 [*pcpA*], and SP_0914) that were enriched in the presence of specific antibiotic mechanisms. Unsurprisingly, *rpoB* mutations, which confer resistance to rifampicin in *S. pneumoniae* and other bacteria^27^, were restricted to rifampicin-treated lineages. More notably, mutations in SP_2040 were enriched in cell wall synthesis inhibitors, and recent evidence shows this gene encodes cell elongation regulator Jag/EloR and interacts with PBP2b, suggesting a novel association and mechanism of tolerance^28^. Convergent mutations in metabolic genes such as *lctO* (lactate oxidase) across multiple drug classes were also detected.

### Validation of Immune-Specific Selection: The Role of Nicotinamidase (SP_1583)

The gene that was most frequently mutated was SP_1583, encoding nicotinamidase, and these nearly exclusively rose to high frequencies in hosts producing neutrophils (i.e., immunocompetent and macrophage-depleted hosts) (**Figure 1D**). This pattern, combined with the fact that all were predicted loss-of-function mutations (frameshift or nonsense substitutions), suggested that loss of nicotinamidase activity confers a specific advantage in the presence of neutrophils. We therefore used a deletion mutant (ΔSP_1583) to test its fitness effects in the presence or absence of neutrophils and observed significant competitive advantages in macrophage-depleted and non-depleted mice (**Figure 1F**). Mechanistically, this advantage appears to be driven by altered cytotoxicity: whereas wild-type TIGR4 induced significant neutrophil death as measured by LDH release, the ΔSP_1583 mutant induced significantly less cytotoxicity, comparable to uninfected controls (**Figure S4B**). This reduction in neutrophil killing was not due to defects in bacterial growth or pneumolysin production in broth culture, nor alterations in neutrophil recruitment, apoptosis, or NETosis *in vivo* (**Figure S4C-H**). Instead, the data indicate that the loss of nicotinamidase production by SP_1583 allows the bacterium to modulate neutrophil cytotoxicity and avoid triggering a damaging inflammatory response. Importantly, this finding validates the power of experimental evolution *in vivo* to identify bona fide adaptations to different immune conditions.

### Host Fitness Costs Select for Tolerance Over Resistance

To evaluate the phenotypic status of the evolved populations we measured levels of drug resistance as minimum inhibitory concentration (MIC) for each endpoint population. Despite hundreds of generations of growth under drug treatments, evolved populations showed minimal to no increase in resistance, with the notable exception of the rifampicin lines (**Figure 2A**). This paradoxical observation, where bacterial populations survived under escalating antibiotic doses without increasing resistance, pointed toward antibiotic tolerance as a potential mechanism. We quantified tolerance by measuring bacterial survival under high-dose antibiotic exposure. Strikingly, 25% (23/90) of the screened lineages exhibited a significant increase in tolerance compared to the ancestral TIGR4 strain, a phenotype that emerged independently of the host immune state (**Figure 2B, S5**).

**Figure 2.**
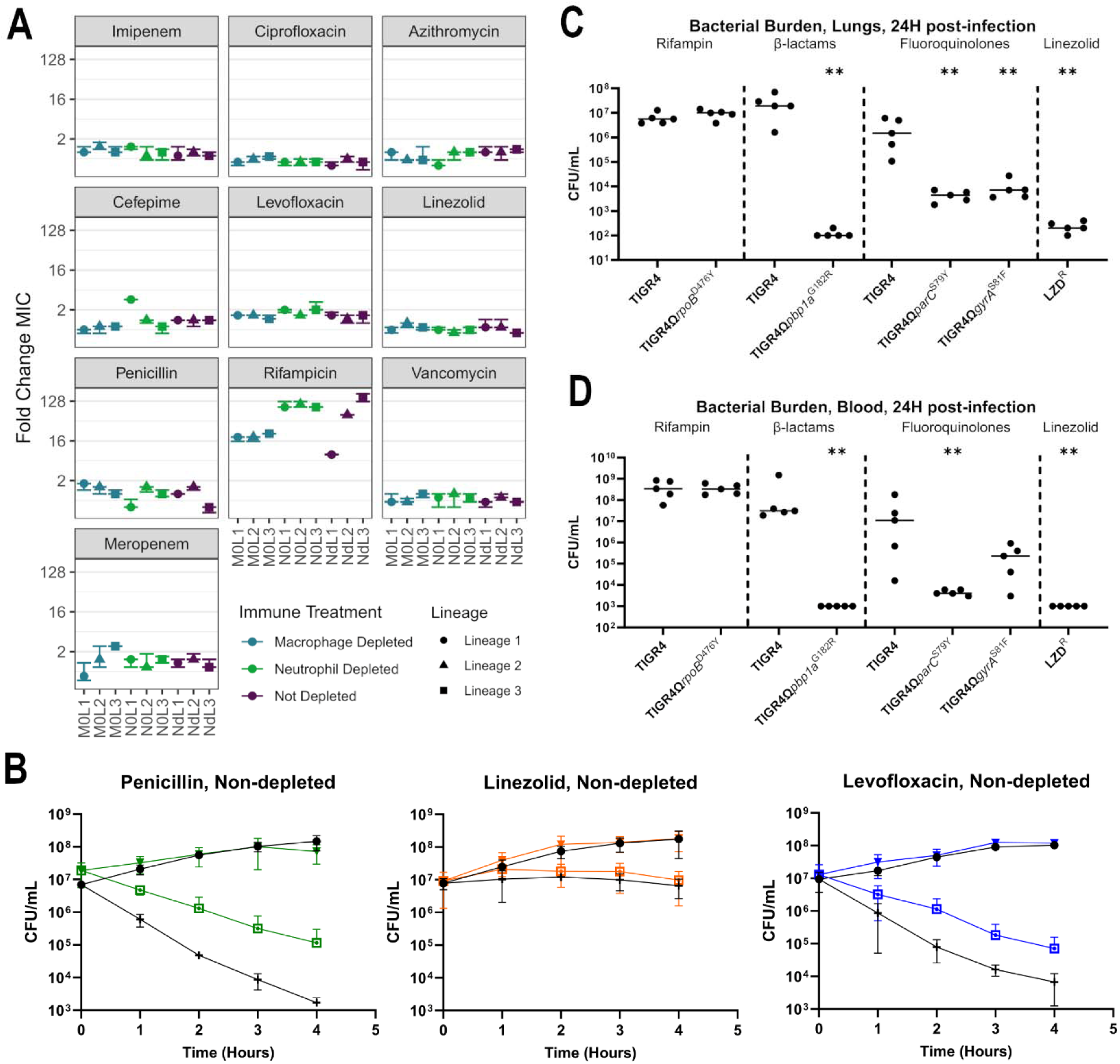
Resistance and fitness characteristics of evolved *S. pneumoniae* populations and mutants. (A) Fold-change in MIC of the final *in vivo*-evolved pneumococcal populations. (B) Antibiotic killing over time in representative populations passaged *in vivo* under penicillin, linezolid, and levofloxacin selective pressures, respectively. Colored lines indicate the evolved population while black lines indicate the original TIGR4 ancestral strain. Punctuated open squares and crosses represent the drug-exposed kill curves for evolved and ancestral strains, respectively. Triangles and circles symbolize the untreated growth curves for evolved and ancestral strains, respectively. (C) Bacterial burden found in the lungs and (D) blood of mice infected with either TIGR4 or antibiotic-resistant mutants of the indicated drugs and drug classes.

We hypothesized that genuine resistance mutations were constrained by costly fitness tradeoffs *in vivo*, limiting their emergence. To test this model, we generated point mutations known to produce resistance to different classes of antibiotics (**Figure 2**) and measured their competitive fitness and ability to cause disease in wild-type mice under drug treatment. These mutations comprised three groups: i) those that emerged during passage but were generally eventually outcompeted (*pbp1a* encoding penicillin-binding protein), ii) those that were never detected (*gyrA* and *parC,* the established targets of fluoroquinolones, and a linezolid resistance mutation) and iii) those that eventually fixed (*rpoB* in the presence of rifampicin). While *rpoB* mutations achieved high population sizes and retained full virulence, all other mutations attenuated the bacteria in the murine pneumonia model (**Figure 2C, 2D**). Because these mutations failed to dominate even in immune-depleted hosts during the evolution experiment despite their certain occurrence based on mutation probabilities^1,21^, it is likely that the physiological cost of these resistance mutations renders them maladaptive in the host. Consequently, *S. pneumoniae* populations may be evolutionarily channeled toward tolerance as a lower-cost strategy for resilience under dynamic antibiotic treatment.

### Convergence on RNase Y: A General Mechanism for Antibiotic Survival

In stark contrast to the loss-of-function mutations found in SP_1583, the next most commonly mutated gene SP_1739 (encoding the RNA degradosome scaffold RNase Y) acquired exclusively single nucleotide polymorphisms (SNPs) across disparate antibiotic classes and immune states (**Figure 1B,D**). The absence of frameshift or nonsense mutations suggests that these variants confer an altered function rather than a simple loss of activity, and further, that these loss of function mutations are deleterious for fitness. Given its convergent evolution across treatments, we hypothesized that RNase Y may confer more generalized resilience under a wide variety of stresses. To explore this model, we first competed an RNase Y deletion mutant (Δ*rny*) against the wild-type strain in the murine lung. In the absence of antibiotics, the Δ*rny* mutant was outcompeted by the wild-type, confirming that RNase Y function is critical for maximal fitness in the absence of stress. However, antibiotic pressure (cefepime or levofloxacin) significantly shifted this dynamic: the wild-type’s competitive advantage eroded, and in the high-stress compartment of the murine lung the Δ*rny* mutant recovered to fitness parity (**Figure 1E)**. This relative fitness gain under severe stress suggests that while total loss of RNase Y is costly, modulation of its activity via the evolved SNPs may provide a survival advantage specifically in the presence of antibiotics.

### Evolved *rny* Variants Confer Broad, Cross-Class Antibiotic Tolerance

To define the causal role of *rny* in antibiotic resilience, we engineered isogenic strains reconstructing eight distinct single nucleotide polymorphisms (SNPs) that reached high frequency in our evolved populations. These included variants selected under fluoroquinolone pressure (A46P, A75G, L77S, R488G, P493R) (**Figure S1F**), β-lactam pressure (V475M, selected by cefepime), and oxazolidinone pressure (A46S, selected by linezolid). Consistent with the phenotypes of the mixed evolved populations, none of these isogenic *rny* mutations significantly elevated the MIC against their respective selection agents (**Figure S6**). However, time-kill assays revealed a striking gain in tolerance to these drugs. Six of the eight variants conferred >1-log enhanced survival against levofloxacin (2x MIC) compared to the parental TIGR4 strain (**Figure 3, S7**). Crucially, this tolerance extended beyond the drug class used for selection. For instance, variants evolved under fluoroquinolone or linezolid pressure (A46S, A46P, R488G, P493R) conferred significant cross-tolerance to the β-lactam cefepime. Similarly, fluoroquinolone-selected variants (A75G, L77S) and the cefepime-selected V475M variant exhibited multi-drug tolerance against both cefepime and the glycopeptide vancomycin. While the precise mutations that evolved experimentally are rare in clinical strains, allelic variants in clinical strains map to similar domains (**Figure S8**). This broad mode of tolerance, where a mutation selected by one antibiotic protects against mechanistically distinct classes, indicates that *S. pneumoniae* has not evolved a specific countermeasure to a single drug target. Instead, these *rny* alleles unlock a generalized stress tolerance program that is accessible and advantageous across diverse antibiotic regimes during murine pneumonia.

**Figure 3.**
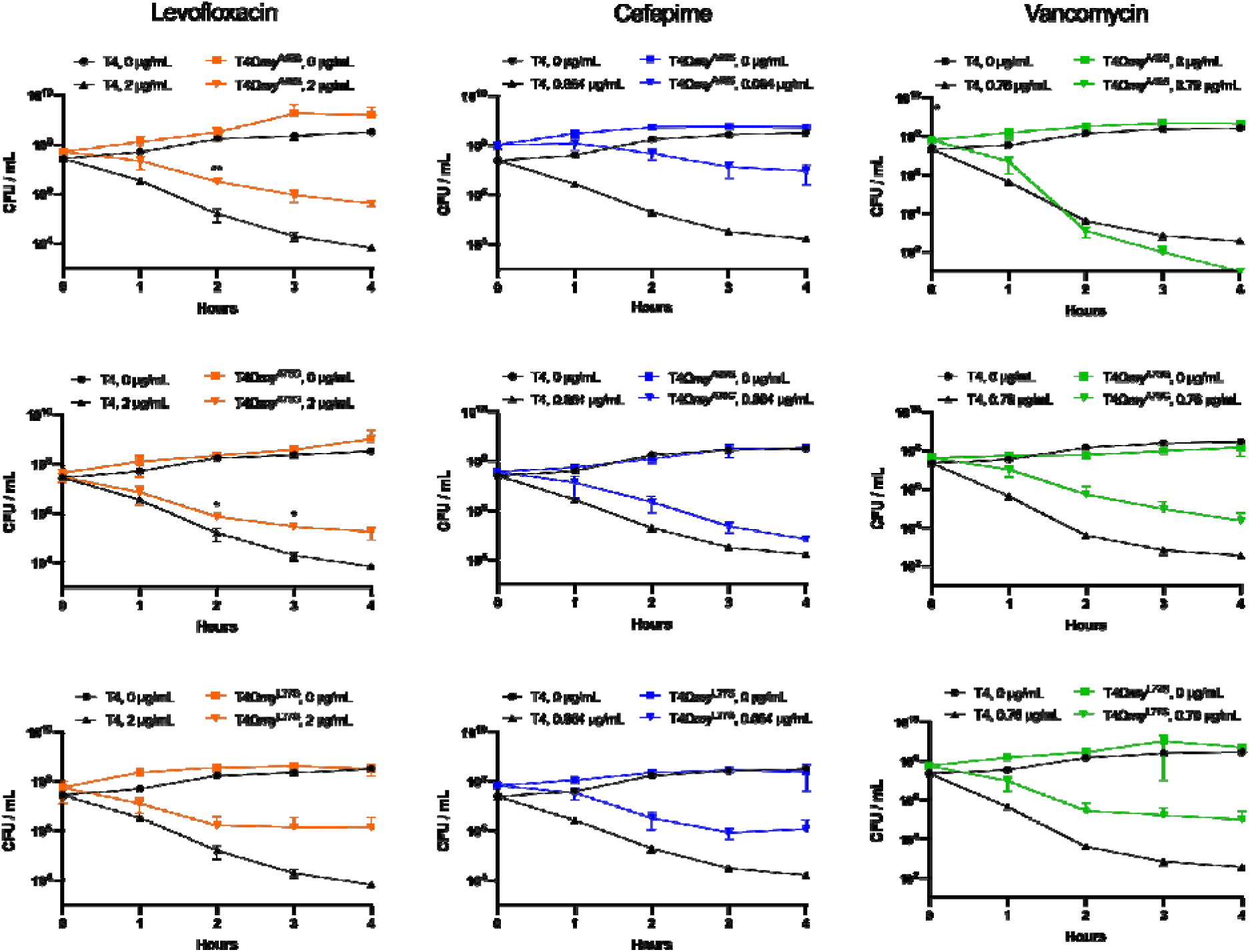
Increased tolerance of evolved RNase Y mutants across drug classes. Black and colored lines represent TIGR4 and *rny* mutants, respectively. Mutant strains were grown to an OD_600_ = 0.2 and exposed to 2× drug MIC values as measured by E-test drug diffusion strips over 4 hours. Colored lines indicate the evolved population while black lines indicate the original TIGR4 ancestral strain. Triangles represent the drug-exposed kill curves for evolved and ancestral strains. Squares and circles symbolize the untreated growth curves for evolved and ancestral strains, respectively.

### Antibiotic Exposure Induces Rapid RNA Clearance and Transcriptional Preservation in ***rny* Mutants.**

As the scaffold for the RNA degradosome, RNAse Y is known to participate broadly in mRNA turnover with varied consequences including stress tolerance, which we and others showed recently^29-31^. To explore how these *rny* mutations affect transcriptional dynamics associated with the observed tolerance, we performed temporal single-cell RNA-Seq (scRNA-Seq) on wild-type (WT) and *rny*L77S populations during exposure to cefepime (2x MIC) and subsequent recovery from treatment (**Figure 4A**). Dimensionality reduction reveals a profound divergence in response strategies to antibiotic exposure. In the absence of antibiotic (T0), both strains occupy a similar transcriptional space (**Figure 4B, 4C**). However, upon antibiotic stress, WT cells diverge wildly from this baseline, forming distinct, disordered clusters that fail to return to the homeostatic state even after antibiotic washout. In contrast, *rny*L77S cells maintain transcriptional similarity to the T0 state across exposure and recovery, with T0 baseline-like profiles becoming increasingly evident during the recovery phase (**Figure 4B, 4C**). This suggests that the *rny* mutation prevents the transcriptional disorder observed in the WT, locking the bacteria into a stable state that is resilient to environmental fluctuations.

**Figure 4.**
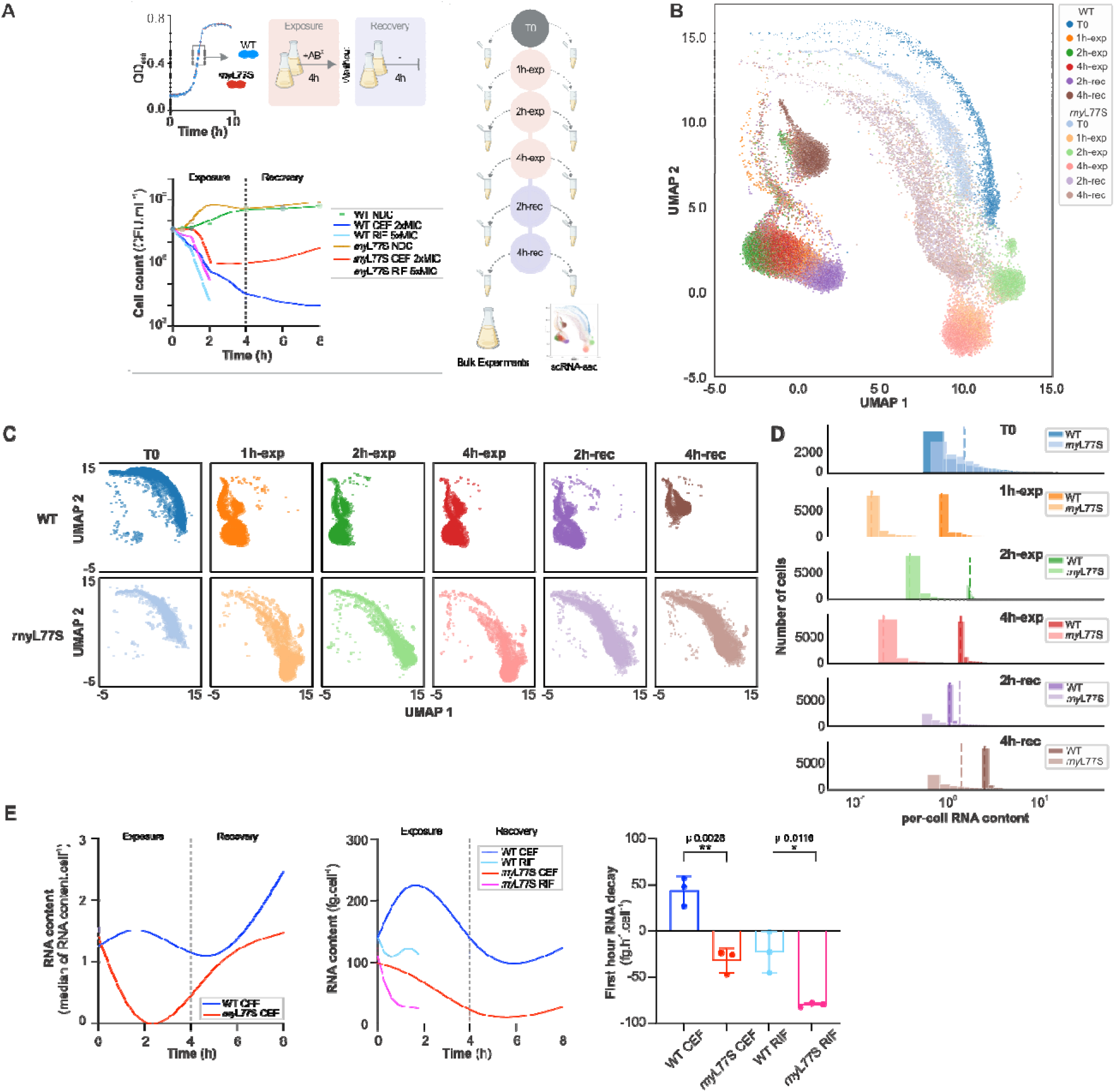
Temporal single-cell RNA-Seq reveals increased RNA decay in *rny*L77S and retention of a baseline-like expression profile. **(A)** Design of temporal single-cell (sc)RNA-seq, bulk RNA collections and corresponding CFU counts during and following drug exposure. The upper left box shows the growth curves of both WT and *rny*L77S in the absence of antibiotics, showing no difference in growth. The bottom left box displays cell counts in the absence and presence of Cefepime (CEF) at 2xMIC, or Rifampicin (RIF) at 5xMIC. *rny*L77S shows increased tolerance to both antibiotics compared to WT. Additionally, *rny*L77S is able to recover faster after CEF washout. The right box shows the temporal sample collection for bulk experiments and scRNAseq during drug exposure and survival. **(B)** UMAP visualization of WT and *rny*L77S single-cell transcriptomes of 10,000 cells at six time points each, colored by sample. **(C)** UMAP plots of the individual single-cell transcriptomes for each strain corresponding to those shown in (**B)**. These individual plots highlight how *rny*L77S retains a cell distribution profile that resembles the unperturbed state, while WT develops a highly divergent state. **(D)** Distribution of per-cell RNA content in WT and *rny*L77S cells at each time point, indicating how *rny*L77S compared to WT rapidly and consistently reduces its RNA content during exposure, and increases RNA content after stress removal. Dashed lines indicate median values. **(E)** Left: The median values of the per-cell RNA content derived from scRNA-seq for WT and *rny*L77S. Middle: As validation for the scRNA-Seq data total RNA was isolated from WT and *rny*L77S cell cultures (bulk experiments) at 30’, 1 and 2 hrs after RIF exposure at 5xMIC, and 30’, 1, 2 and 4 hrs after CEF exposure at 2xMIC and at 2 and 4 hrs after CEF washout. Right: During the first hour of drug exposure, the rate of decay is significantly faster in *rny*L77S, both in the presence of CEF or RIF.

### Mutations in the RNase Y Scaffold Drive Hyper-Active RNA Turnover

We hypothesized that the transcriptional stability of *rny* mutants stems from altered kinetics of mRNA decay. RNase Y is a critical scaffold of the bacterial degradosome; pneumococcal RNase Y shares 76% amino acid similarity with its *B. subtilis* homolog, where the N-terminal domain anchors the protein to the membrane and additional domains recruit enzymatic partners like RNase J, RppH, and enolase^32-34^. Notably, most evolved mutations, including L77S, cluster within the N-terminal unstructured scaffold domain or the C-terminus (**Figure S1F**). We postulated that these structural changes alter the recruitment or efficiency of the degradosome.

To investigate this dynamic, we first determined that scRNA-Seq data can be used to estimate the amount of RNA per cell, by summing normalized expression counts per cell followed by normalization to a sequencing depth of one million reads per sample. Using this metric, we found that upon CEF exposure *rny*L77S experiences a pronounced and rapid decline (∼10-fold) in total RNA content (**Figure 4D**). In contrast, while the distribution of RNA per cell for WT overall becomes tighter, its cells retain an amount of RNA that is much closer to the unperturbed state and significantly more than L77S during CEF exposure (Mann–Whitney□U□test,□p□<□7.6□×□10□^23^), **Figure 4D**). We validated this phenomenon using bulk RNA quantification, which captures total RNA including rRNA that are excluded in our scRNA-Seq analyses. The bulk kinetics confirmed a distinct bipartite response: upon cefepime exposure, WT cells initially increase total RNA, likely reflecting an abortive (r)RNA transcriptional surge, before degrading it only in the late exposure phase (2–4 hours) (**Figure 4E**). Conversely, *rny*L77S bypasses this accumulation entirely, immediately initiating rapid RNA clearance. To isolate degradation from synthesis, we quantified decay rates in the presence of rifampicin (an RNA synthesis inhibitor). The *rny*L77S mutant exhibited an RNA decay rate at least 3.5-fold higher than the WT during the first hour of stress (unpaired t-test, p = 0.0116; **Figure 4E**). This confirms that while the mutant may still attempt active transcription, its efficient degradation machinery overwhelms synthesis, enforcing a low-RNA state. Crucially, this state is reversible: upon antibiotic removal, both strains resume RNA synthesis. However, while *rny*L77S successfully reconstructs its original homeostatic profile, the WT only partially recovers RNA quantity and fails to overcome the perturbed transcriptional state.

### Single-Cell Clustering Reveals a Dual Survival Strategy

To deconstruct the population-level heterogeneity driving *rny*-mediated tolerance, we applied unsupervised Louvain clustering, which identifies non-overlapping communities within large networks (resolution 0.1), to map distinct transcriptional states across the exposure and recovery timeline (**Figure 5A**). This analysis resolved four primary clusters that clearly differentiate the wild-type and mutant responses. The wild-type population is confined to Clusters 1 and 4. These cells, present during both cefepime exposure and the recovery phase, are characterized by high transcriptional entropy, a metric we have previously established as a proxy for systemic cellular disorder and impending cellular death^35^. Even after antibiotic removal, WT cells in Cluster 4 fail to restore order, retaining high entropy levels comparable to the stress state (**Figure 5D, S9A**). In contrast, the *rny*L77S population bifurcates into two distinct lineages upon stress exposure: I. Cluster 2, a quiescent majority, which dominates during antibiotic exposure. It is characterized by significantly reduced RNA content (**Figure 5C**) and elevated entropy, reflecting a transcriptionally dormant state; II. Cluster 3, a resilient minority, comprising ∼12.2% of the population after 1 hour of exposure, these cells are transcriptionally indistinguishable from the unperturbed baseline (T0) and recovery populations (**Figure 5A, 5B**). Crucially, Cluster 3 maintains consistently low entropy throughout the experiment (**Figure 5D**), suggesting these cells successfully buffer the antibiotic stress to maintain homeostasis.

**Figure 5.**
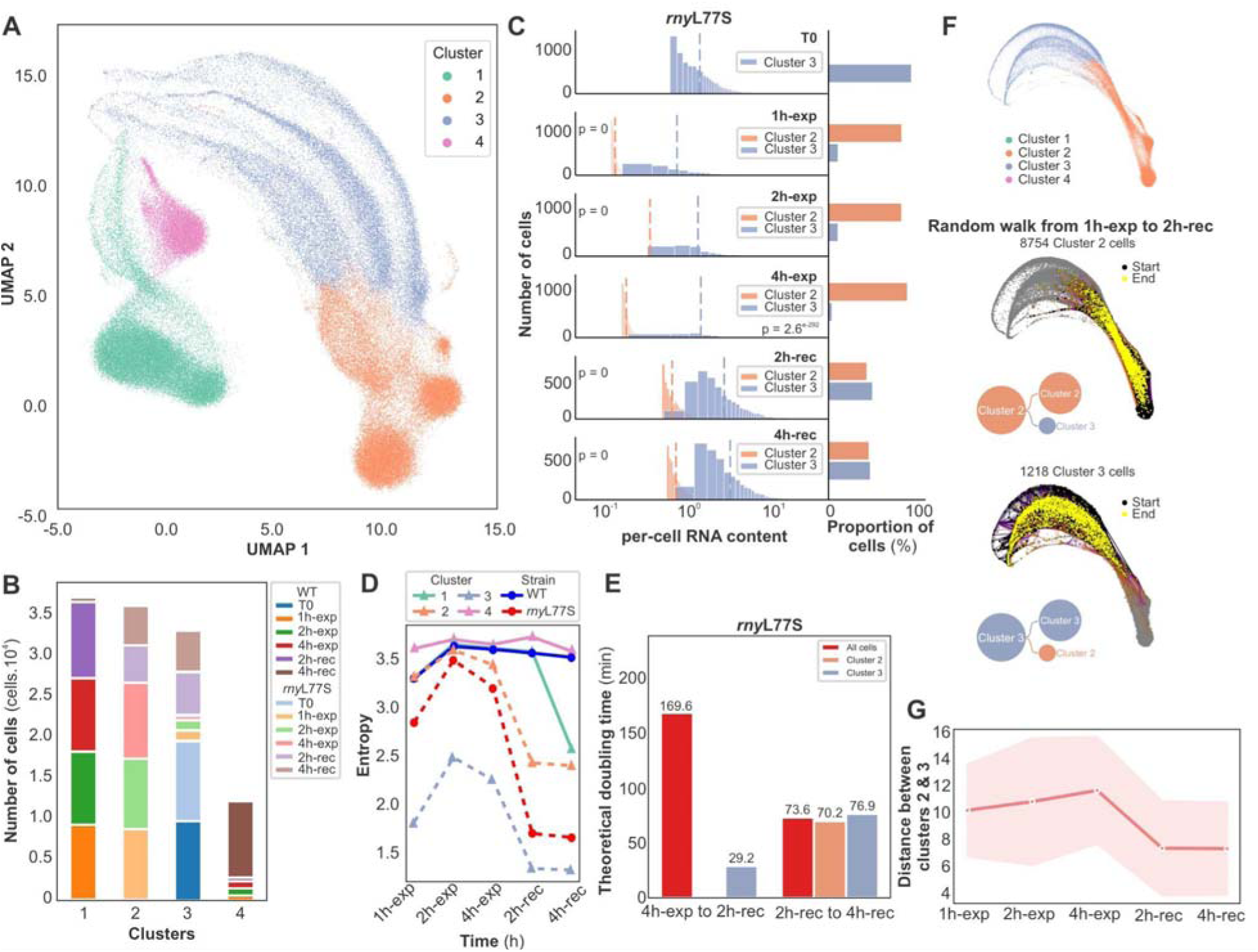
Single-cell transcriptomic clustering reveals dual survival strategies under antibiotic stress. **(A)** Louvain clustering of the 12 temporal single-cell WT and *rny*L77S profiles at resolution 0.1, colored by clusters. **(B)** Cell counts per sample for each cluster shown in (**A)**, highlight how clusters 1 and 4 consist of WT cells, cluster 2 of *rny*L77S drug exposure and recovery cells, and cluster 3 mainly of unperturbed and *rny*L77S recovery cells. **(C)** Distribution of per-cell RNA content for *rny*L77S cells (left, histogram) and the percentage of cells (right, bar plot) in Clusters 2 and 3 at each time point, which highlights how cluster 2 cells with low RNA content dominate during CEF exposure, and cells shift towards cluster 3 with increased RNA content during recovery. Dashed lines indicate median RNA content. *P*-values from two-sided Mann–Whitney U tests comparing clusters 2 and 3 are shown in the plot. **(D)** Entropy at each time point compared to T0 within each cluster in WT (marked with blue circle) and *rny*L77S (marked with red circle). Clusters dominated with WT cells are shown as solid lines, while clusters with *rny*L77S cells are shown as dashed lines. *rny*L77S cells have lower entropy than WT, indicative of their enhanced ability to overcome CEF exposure. Moreover, cluster 3 cells retain a low entropic state throughout, while cluster 2 cells can rapidly reduce entropy after stress removal indicative of their ability to contribute to recovery. Cluster–timepoint combinations with fewer than 250 cells are excluded from the plot. **(E)** Theoretical doubling time estimated from clusters 2 and 3 and for all *rny*L77S cells between 2h recovery (2h-rec) and 4h exposure (4h-exp), and between 2h recovery (2h-rec) and 4h recovery (4h-rec). If cells in cluster 3 are solely responsible for recovery their doubling time would be ∼29’ in the first 2 recovery hours, while it would drop to ∼76’ from 2 to 4 h. (**F**) Top: Force-directed layout of pooled *rny*L77S cluster 2 and cluster 3 cells. 10,000 CellRank random walk simulations from 1 h exposure (1h-exp) as start point (black dots) to 2 h recovery (2h-rec) as end point (yellow dots) show that while most cells remain within their original (starting) cluster, a subpopulation of cells from cluster 2 (middle) and cluster 3 (bottom) are capable of transitioning from one cluster to the other. This indicates that the transcriptional similarity between cluster 2 and cluster 3 cells supports, at least theoretically, that cluster 2 cells are able to change their profile and contribute to recovery. (**G**) Based on transcriptional similarities (measured as the distribution of Euclidean distances per-cell between cluster 2 and 3) cells from cluster 2 and 3 initially diverge from each other during CEF exposure, but then converge during recovery, again indicating cluster 2 cells are changing into cluster 3 cells to contribute to recovery. Solid line represents the mean, and the shaded area indicates the standard deviation

### Reactivation of the Quiescent Majority Contributes to Population Recovery

The existence of these two subpopulations raised a critical question of whether post-antibiotic recovery is driven solely by the division of the resilient Cluster 3 or if the quiescent Cluster 2 contributes to reactivation. We addressed this quantitatively by analyzing population dynamics over the first 2 hours of recovery (**Figure 5E**). During this window, the proportion of Cluster 3 cells expanded from ∼10% to ∼50% (**Figure 5B**), while the total population only doubled (total doubling time ∼169 minutes). If this population growth was fueled exclusively by the division of the resilient Cluster 3 survivors, these cells must double every ∼29 minutes to account for the observed biomass increase. This rate is biologically implausible, as it exceeds the maximum growth rate of *S. pneumoniae* under optimal unperturbed conditions (∼35–40 minutes). Furthermore, this model would imply a sudden deceleration to a 76-minute doubling time in the subsequent window (2–4 hours), which is inconsistent with exponential recovery.

Therefore, the data necessitate a model where recovery is driven cooperatively: the resilient Cluster 3 divides, while the quiescent Cluster 2 exits dormancy and transitions into the healthy Cluster 3 state. This reactivation model is supported by CellRank time-course analysis^36^, which estimates the likelihood of cellular transitions from one cluster to another across time points. Random-walk simulations indicate a high probability of cells transitioning from Cluster 2 to Cluster 3 during the recovery phase (**Figure 5F, S9B**). Additionally, Euclidean distance analysis confirms that while the clusters diverge transcriptionally during stress, they rapidly reconverge upon drug removal, indicating that Cluster 2 cells are transcriptionally reprogrammed to match the homeostatic Cluster 3 profile (**Figure 5G**). Collectively, these findings demonstrate that *rny* mutants employ a bet-hedging strategy: a minority subpopulation maintains a near baseline transcriptional profile to allow immediate outgrowth, while the majority enters a reversible, low-RNA quiescent state to tolerate and subsequently facilitate regrowth in response to dynamic antibiotic exposure.

### High-Fidelity RNA Preservation and Ribosomal Turnover Drive Tolerance

Given that wild-type (WT) cells retain higher total RNA content yet fail to survive, we investigated the integrity of the retained transcriptome. We observed that WT RNA quality degrades progressively throughout cefepime exposure and fails to recover even after drug removal (**Figure 6A, S10**). In striking contrast, *rny*L77S cells, despite their rapid bulk RNA loss, maintain significantly higher RNA quality throughout both exposure and recovery (**Figure 6B**). This suggests that the *rny* mutant actively purges damaged or unnecessary transcripts to preserve a high-fidelity core transcriptome, preventing the accumulation of toxic, fragmented RNA that characterizes the WT programmed death. To define the selectivity of this purge, we calculated normalized degradation rates for each transcript in each sample. We found that *rny*L77S exhibits a global decay rate ∼6-fold faster than WT (5.64 ± 1.2 vs. baseline; Supplementary File 1), with a specific bias towards degradation of the most abundant transcripts (**Figure 6B; S11A-F**). We found this degradation is heavily skewed toward translation-related genes, which coincides with the preferred degradation of essential and 30S and 50S ribosomal protein transcripts (Mann–Whitney U test, p < 7.37 × 10□□; **Figure 6C, 6D, S11G**).

**Figure 6.**
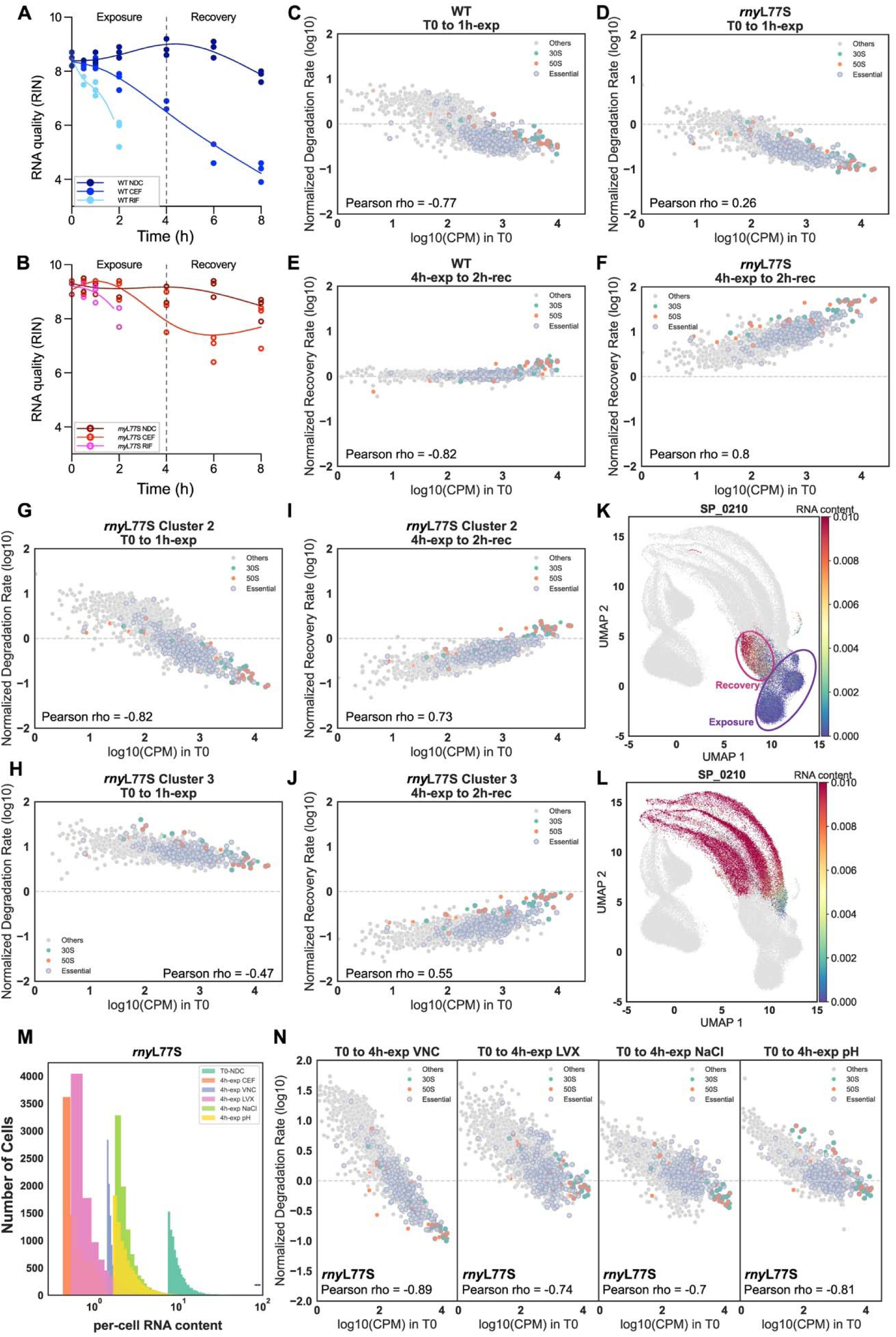
Preservation of high quality RNA and enhanced decay and recovery of ribosomal transcripts drives a general tolerance response in *rny*L77S. **(A–B)** RNA Integrity Number (RIN) scores for (**A**) WT and (**B**) *rny*L77S cells in the absence or presence of CEF, or RIF for up to 4 h, and after 2 h and 4 h of recovery following drug washout (total RNA was used from ‘bulk’ experiments in **Fig.1E** middle panel). *rny*L77S cells maintain higher RNA quality during antibiotic exposure compared to WT cells, under both CEF and RIF treatment, as well as after CEF removal. **(C-D)** Normalized degradation rates from T0 to 1 h exposure (1h-exp) plotted against log10 counts per million (CPM) for WT (**C**) and *rny*L77S (**D**). This shows that *rny*L77S transcripts degrade faster than WT transcripts, with preferred degradation of essential and 30S and 50S ribosomal protein transcripts. **(E-F)** Normalized recovery rates from 4h exposure (4h-exp) to 2h recovery (2h-rec) plotted against the log10 CPM in T0 for WT (**E**) and *rny*L77S (**F**). During recovery, while all transcripts follow a similar recovery rate in WT, global transcription and especially essential and 30S and 50S ribosomal protein transcripts recover faster in *rny*L77S. **(G-H)** Normalized degradation rates from T0 to 1h-exp are plotted against the log10 CPM in T0 for *rny*L77S cluster 2 (**G**) and cluster 3 (**H**) cells. These results show that cluster 2 cells have a faster RNA degradation rate, with essential, and 30S and 50S ribosomal protein transcripts preferentially degraded. Conversely, RNA degrades at a slower rate in cluster 3 cells compared to the entire *rny*L77S population. **(I-J)** Normalized recovery rates from 4h-exp to 2h-rec are plotted against the log10 CPM in T0 for *rny*L77S cluster 2 (**I**) and cluster 3 (**J**) cells. These results show that transcripts for essential and 30S-50S ribosomal proteins recover faster than the other transcripts in both clusters. Note that all rates are normalized to WT (**C-F**) or rnyL77S (**G-J**) to enable direct comparisons. **(K-L)** UMAP in which cluster 2 (**K)** and cluster 3 (**L**) cells are colored by RNA content for SP_0210/*rplD*, which shows that RNA content is uniformly maintained within cluster 3, but varies in cluster 2 with a lower content during CEF exposure. Notably, SP_0210 RNA content increases in cluster 2 cells during recovery, reaching values similar to those of cluster 3 cells, indicative of cluster 2 cells resembling cluster 3 cells over time and contributing to recovery. Blue oval highlights *rny*L77S cells from recovery samples (2h/4h-rec); red oval highlights *rny*L77S cells from exposure samples (2h/4h-exp). **(M)** Distribution of per-cell RNA content at T0, and 4h exposure to CEF, vancomycin (VNC), levofloxacin (LXV), high osmolarity (300 mM NaCl), and acidic conditions (pH 4.2), confirming rapid global RNA decay as a general tolerance response in *rny*L77S. **(N)** Normalized degradation rates from T0 to 4h-exp plotted against the log10 CPM in T0 for the same conditions as in (**M)**, highlighting similar patterns as those found under CEF exposure, with essential and 30S-50S ribosomal protein transcripts preferentially degraded. All Pearson correlation P values in the related plots are < 4.26× 10^□5^.

### A "Ribosomal Reboot" Strategy Facilitates Rapid Recovery

Importantly, this selective degradation sets the stage for the accelerated recovery of *rny*L77S. Upon antibiotic removal, the WT attempts a generalized transcriptional restart, regenerating all gene classes at similar rates. The *rny*L77S mutant, however, executes a prioritized "ribosomal reboot" whereby transcripts encoding essential genes and ribosomal proteins are regenerated significantly faster than the rest of the transcriptome (Mann–Whitney U test, p < 4.02 × 10□²□; **Figure 6E, 6F, S11H**). This suggests the mutant’s strategy is to strip down to a minimal state and then immediately prioritize the reconstruction of the translational machinery to drive rapid growth in response to fluctuating antibiotic exposure. We confirmed that this strategy is utilized by both the resilient (Cluster 3) and quiescent (Cluster 2) subpopulations identified in our clustering analysis. While Cluster 2 cells show more dramatic dynamics, transitioning from profound RNA depletion during exposure to a massive transcriptional rebound during recovery (i.e., faster transcript degradation and recovery rates compared to Cluster 3; Mann–Whitney U test,□p□=0, **Figure 6G-J**), both clusters exhibit the same statistical preference for recovering ribosomal transcripts (Mann–Whitney U test, p < 1.23 × 10□¹□; **Figure 6G-J, S12A-F**). Single-cell analysis of representative ribosomal genes (e.g., *rplD* / SP_0210) highlights this convergence with the Cluster 2 cells, which suppress *rplD* during stress, rapidly ramping up expression during recovery to match the levels of Cluster 3 (**Figure 6K, 6L, S12G-P**). This confirms that the quiescent majority (Cluster 2) is successfully reprogrammed to rejoin the active population via this ribosomal recovery program. These findings highlight that ribosomal RNA turnover is a key determinant of stress resilience, underpinning a dual survival strategy in *rny*L77S.

### The *rny*-Mediated Stress Response is Universal

Finally, we asked if this mechanism explains the broad cross-tolerance observed earlier. We exposed *rny*L77S cells to a panel of distinct stressors, including vancomycin, levofloxacin, hyperosmolarity (300 mM NaCl), and acidity (pH 4.2). Importantly, in all conditions, the mutant exhibited enhanced survival. scRNA-Seq profiling confirmed that the underlying mechanism is conserved with the rapid global RNA decay coupled with the preferential degradation of ribosomal and essential transcripts (Mann–Whitney U test, p < 4.82 × 10□□; **Figure 6M, 6N, S13**).

Besides increased survival this transcriptional reset mechanism has a direct macroscopic consequence: accelerated recovery following antibiotic insult. Since lag-phase duration is a critical determinant of survival in fluctuating *in vivo* environments^37,38^, we measured the time required for evolved mutants to resume growth after antibiotic removal. We utilized shortened antibiotic exposures whereby the parental and mutant strains produced equivalent numbers of viable cells prior to antibiotic removal. Strikingly, *rny* mutants consistently exited lag phase several hours earlier than the parental strain (**Figure 7**). This phenotype was independent of the antibiotic mechanism of action, confirming that modulation of RNase Y unlocks a generalist survival program that minimizes the refractory period following population collapse.

**Figure 7.**
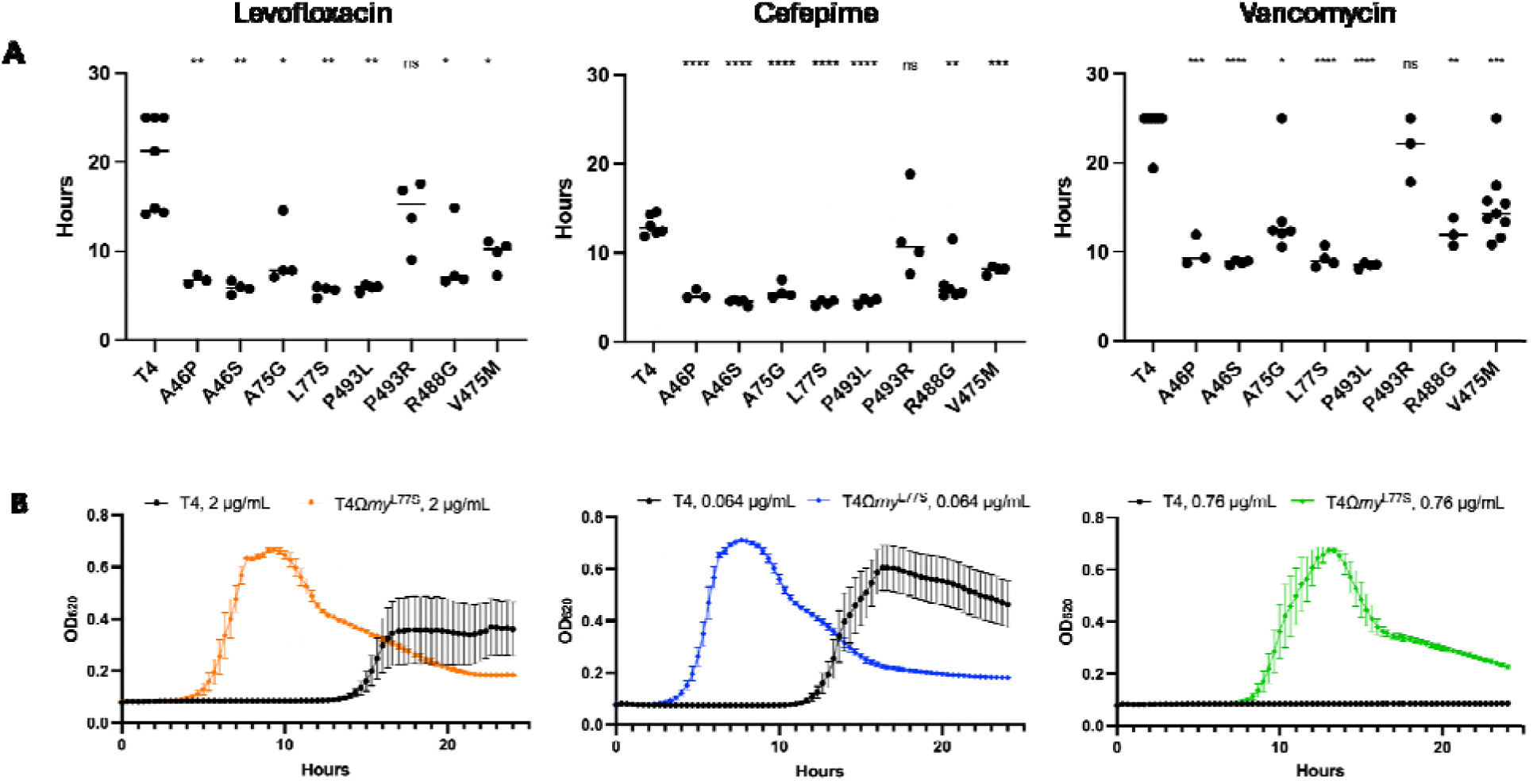
Mutations in *rny* Facilitate Reduced Lag-phase and Rapid Recovery Following Antibiotic exposure. **A)** The parental wildtype (T4) or the respective point mutants in *rny* were subjected to transient antibiotic exposure followed by back-dilution and monitoring for resumption of bacterial growth. In response to different classes of the antibiotics, the *rny* mutants demonstrated significantly reduced lag phase following antibiotic removal (A). Each data point indicates an independent biological replicate, significance determined by ANOVA, *=p<0.05, ** p<0.01, *** p<0.005, **** p<0.0001. **B)** Growth curves show shorter duration of lag-phase for the *rny* L77S mutant in comparison to the parental T4 following antibiotic exposure. Data represent the average and standard deviation from at least three biological replicates.

## DISCUSSION

### The Host Environment Imposes Selective Constraints that Suppress the Emergence of Resistance

Elucidating the evolutionary dynamics at the intersection of antibiotic pressure and host immunity is critical for understanding treatment failure. While prolonged antibiotic exposure in laboratory settings reliably drives the emergence of high-level resistance^15,39,40^, our large scale *in vivo* analysis demonstrates that the host environment imposes stringent selective constraints that fundamentally redirect adaptive trajectories. This demonstrates the power of experimental evolution *in vivo*, which enables a nuanced forward genetic screen for mutations that balance microbial fitness in the host with the ability to endure a given stressor, in this case antibiotics. We observed that *S. pneumoniae* populations evolved in the murine lung rarely acquired canonical resistance mutations, even under escalating antibiotic dosage. The notable exception was rifampicin, where *rpoB* mutations confer high-level resistance with minimal physiological cost^41,42^. For all other antibiotic classes, the fitness costs associated with frank resistance appear prohibitive in the context of infection. Our competitive assays confirmed that resistance alleles capable of withstanding antibiotic pressure *in vitro* rendered the bacteria competitively inferior *in vivo*. This supports a model where the requirement to survive immune clearance exerts purifying selection against high-cost resistance mutations, effectively channeling the population toward alternative survival strategies^37^. For example, convergent mutations in metabolic genes such as *lctO* (lactate oxidase) were observed across multiple drug classes. Since L-lactate can function as a hormetic promoter of oxidative stress resistance^43^, *lctO* mutations likely represent a generalized metabolic adaptation to antibiotic-induced oxidative stress rather than a drug-specific resistance mechanism.

### Convergence on *rny*: A Generalist Strategy for Stress Tolerance

Deprived of classical resistance pathways, pneumococcal populations convergently evolved mutations in *rny*, encoding the RNA degradosome scaffold RNase Y. Unlike drug-specific resistance mechanisms, *rny* mutations emerged across diverse antibiotic classes and immune states, conferring a broad, cross-protection phenotype. This suggests that modulation of RNA turnover represents a generalist solution to environmental stress. By altering the kinetics of the degradosome, these mutations allow the bacterium to bypass the specific mechanism of action of the drug and instead survive the downstream physiological crisis it induces. This mirrors findings in *Mycobacterium*, where alteration of RNase J confers multidrug tolerance^44^, and highlights the central role of RNA metabolism in the intrinsic resistance of Gram-positive pathogens^45-50^.

### Mechanism of Survival: Averting Transcriptional Collapse

The distinction between the wild-type (WT) death and *rny* mutant survival reveals a fundamental insight into the nature of antibiotic lethality. We propose that antibiotic-induced death in *S. pneumoniae* is driven by transcriptional collapse, a chaotic surge in transcription followed by a rapid loss in RNA quantity and integrity. Specifically, the WT response is characterized by high entropy^35^, where the cell initially attempts to transcribe its way out of the crisis, but ends up with fragmented, low-quality RNA and eventual systemic failure. In contrast, *rny* mutants avert this collapse by enforcing a state of controlled quiescence. The hyper-active degradation observed in the *rny*L77S mutant effectively clears the transcriptional slate. Rather than allowing damaged transcripts to accumulate, the mutant rapidly degrades the bulk of its RNA. This serves two critical functions, reducing the entropic burden on the cell and preventing the translation of aberrant proteins. Crucially, this degradation is selective; the mutant preserves a core of high-quality transcripts, allowing it to maintain homeostasis in a dormant state.

### The "Ribosomal Reboot" and Bet-Hedging

Importantly, our single-cell analysis reveals a dual bet-hedging strategy. A minority subpopulation (Cluster 3) maintains a near-baseline profile, ready for immediate outgrowth, while the majority (Cluster 2) enters a quiescent state. The key to the mutant’s rapid recovery, the short lag phase phenotype, is its prioritization of the translational machinery. Upon stress removal, the mutant executes a rapid ribosomal recovery, regenerating 30S and 50S transcripts significantly faster than the rest of the genome. This ensures that as soon as conditions are permissive for growth, the cell has the translational capacity to resume exponential growth, a major competitive advantage in the fluctuating environment of the host.

### Immune-Specific Adaptations and Evolutionary Trade-offs

While *rny* represents a generalist adaptation,, our powerful screen using experimental evolution also captured distinct immune-specific trajectories. The convergence on *SP_1583* (nicotinamidase) loss-of-function mutations exclusively in neutrophil-replete hosts highlights the specificity of immune pressure. By modulating nicotinamide metabolism, these mutants likely dampen neutrophil-mediated cytotoxicity^51-53^, identifying a novel route of immune evasion. Similarly, convergent evolution of mutations in SP_2040 encoding the cell elongation regulator Jag/EloR in cell wall synthesis inhibitors suggests a novel mechanism of tolerance, and parallel mutations in metabolic genes such as *lctO* (lactate oxidase) point to a generalized metabolic adaptation to oxidative stress. Finally, if *rny*-mediated tolerance is so advantageous, why has it not become the driving mechanism to overcome stress in the species? We posit that this strategy carries a conditional cost. While the mechanism of aggressive transcript clearance is protective during severe stress it is likely maladaptive under benign or less stressful conditions, where it would necessitate futile cycles of transcription and degradation and hence be energetically expensive. Thus, these mutations represent a classic evolutionary trade-off: a specialized adaptation for high-stress environments that is maintained at low frequencies in the population or selected *de novo* when the drug-host-pathogen interface becomes lethal. Collectively, these data identify RNA turnover not just as a housekeeping function, but as a tunable rheostat for bacterial survival, offering a potent conceptual target for combating antibiotic tolerance.

## ACKNOWLEDGMENTS

This research was supported by NIH grant(s) (5U01AI124302, 7U19AI158076, 7R01AI148470) and the American Lebanese Syrian Associated Charities (ALSAC). We would like to thank Victor Torres and his laboratory trainees Adam Pickrum and Amelia Pi for their guidance in human neutrophil work and help in obtaining and purifying human neutrophils, respectively. We also thank the laboratory of Elaine Tuomanen for gifting the pneumolysin antibody. The content is solely the responsibility of the authors and does not necessarily represent the official views of the National Institutes of Health.

## AUTHOR CONTRIBUTIONS

Conceptualization, T.V.O., R.R.I., V.S.C., J.W.R., A.R.I., A.T.N., M.R.S., J.O.M., and Y.Y.; Methodology, T.V.O., V.S.C., J.W.R., A.R.I., A.T.N., and J.O.M.; Formal Analysis, M.R.S., A.T.N., Y.Y., J.O.M. and Q.J.; Investigation, A.T.N., M.R.S., J.O.M., Y.Y., H.E., A.R.I., A.E.M., A.P., E.M., J.C., N.O., R.K.L., and A.R.; Resources, J.W.R., M.E.H., V.S.C., T.V.O., and J.O.M.; Writing – Original Draft, A.T.N., M.R.S. J.O.M., Y.Y.; Writing – Review & Editing, A.T.N., M.R.S., J.O.M., Y.Y., T.V.O., V.S.C., J.W.R.; Supervision, J.W.R., V.S.C., T.V.O.

## STAR METHODS

### KEY RESOURCES TABLE

### RESOURCE AVAILABILITY

#### Lead contact

Further information and requests for resources and reagents should be directed to and will be fulfilled by the lead contact, Jason Rosch.

#### Materials availability

Streptococcal populations and strains used in this study are readily available upon request from the lead contact.

#### Data and code availability

- All sequence data have been deposited at the NCBI SRA database and are publicly available as of the date of publication. Accession numbers are listed in the key resources.
- All original code used for genome sequencing data analysis, visualization, and statistical analyses of mutations is available on GitHub on GitHub (https://github.com/michellescribner/spn-evo).
- Python scripts and all necessary input files used for single-cell RNA-seq downstream analyses and figure generation are available in Mendeley Data (DOI: 10.17632/6j3ckhtp8x.1).
- Any additional information required to reanalyze the data reported in this paper is available from the lead contact upon request

### EXPERIMENTAL MODEL AND SUBJECT DETAILS

#### Mouse models

Female wildtype BALB/c mice were purchased from The Jackson Laboratory and housed in groups of five under a 12-hour light and dark cycle with regular husbandry handled by in-house staff of the St. Jude Animal Resource Center. All mice were approximately seven weeks old at the outset of all experiments, and mice were randomly assigned to experimental groups. All experiments, protocols, and procedures involving mice were performed with approval of and in accordance with the NIH Guide for the Care and Use of Laboratory Animals and guidelines of the St. Jude Institutional Animal Care and Use Committee.

### METHOD DETAILS

#### Strains and growth conditions

For initial growth, all pneumococcal strains were streaked from 20% glycerol frozen stocks onto tryptic soy agar (Millipore Sigma, Billerica, MA) containing 3% defibrinated sheep’s blood and 20 µg/mL neomycin and grown overnight at 37 °C with 5% CO_2_. For growth in liquid culture, pneumococcal colonies were inoculated into semidefined C+Y media or Todd-Hewitt Broth containing 2% yeast extract ^54^.

#### Pneumococcal mutant generation

To generate the SP_1583 (nicotinamidase) deletion mutant T4Δ1583::PhSE in the pneumococcal TIGR4 strain, we PCR amplified an approximate 1000 bp region containing homology to the immediate upstream and downstream regions flanking the SP_1583 open reading frame (ORF). The 5’ flanking region was PCR amplified using primers SP1583_AB_F and SP1583_AB_R (**Supplementary File 1**). The 3’ flanking region was PCR amplified by SP1583_CD_F and SP1583_CD_R. Primers SP1583_AB_R and SP1583_CD_F additionally contained an approximate 27-30 bp sequence homologous to the 5’ and 3’ ends of the PhunSweetErm selection cassette, a modified version of the previously described PhunSweet cassette, containing an erythromycin resistance marker in place of the kanamycin resistance gene ^55^. Briefly, the PhunSweet cassette without the kanamycin resistance gene was amplified from TIGR4Δcps::PhunSweet gDNA using primers PhunSweet_F and PhunSweet-kan_R to generate fragment PhunSweet-kan (**Supplementary File 1**) ^56^. The erythromycin resistance cassette erm was amplified from TIGR4ΔspxB::Erm gDNA using primers Erm_F and Erm_R to generate fragment Erm. PhunSweet-kan and Erm PCR products were spliced together using PhunSweet_F and Erm_R primers using SOE PCR to form the PhunSweetErm cassette ^57^. Utilizing Primers SP1583_AB_F and SP1583_CD_R, the PhunSweetErm cassette was then fused via three-fragment SOE PCR between the PCR products containing the 5’ and 3’ regions flanking the SP_1583 ORF.

To create point mutations in SP_1739, we first generated a deletion mutant of SP_1739 by amplification and SOE PCR of 1000 bp regions flanking the gene of interest fused to the PhunSweetErm cassette as with the SP_1583 mutant construction. Primers sets SP1739_AB_F & SP1739_AB_R and SP1739_CD_F & SP1739_CD_R were used to amplify the 5’ and 3’ region flanking the SP_1739 ORF, respectively. Successful deletion of the ORF and insertion of the cassette generated strain T4Δ1739::PhSE, which was used as the parent strain for SP_1739 point mutant creation. Point mutations for the A46P, A46S, A75G, L77S, V475M, R488G, P493L, and P493R amino acid changes in SP_1739 were all generated via two fragment SOE PCR (Table S1). The resulting PCR products containing 5’ and 3’ homology to SP_1739 and the mutated ORF were transformed into strain T4Δ1739::PhSE to generated strains T4Δ1739::1739^A46P^, T4Δ1739::1739^A46S^, T4Δ1739::1739^A75G^, T4Δ1739::1739^L77S^, T4Δ1739::1739^V475M^, T4Δ1739::1739^R488G^, T4Δ1739::1739^P493L^, and T4Δ1739::1739^P493R^. Mutations in each strain were confirmed after amplification of the SP_1739 gene and Sanger sequencing using primers SP1739_SeqA through SP1739_SeqH.

#### Antibiotic tolerance and recovery testing

In order to determine whether antibiotics demonstrated a decreased rate of killing against tolerant RNase Y mutant strains, TIGR4 and RNase Y point mutants were grown to an OD_620_ ∼ 0.2 in the pneumococcal logarithmic phase of growth. Antibiotic was added to bacterial liquid cultures at 1×, 2×, or 4× MIC of each tested strain. Bacterial CFUs were enumerated from sample titrations taken from the bacterial culture prior to antibiotic expose (t = 0) and every hour afterwards for 4 hours. In order to measure the length of time the RNase Y mutants took to exit lag phase post antibiotic exposure, TIGR4 and RNase Y point mutants were grown in C+Y media to an OD_620_ ∼ 0.2. Antibiotic was added to bacterial liquid cultures at 2x MIC, corresponding to 2 μg.mL ^-1^ for levofloxacin, 0.76μg.mL ^-1^ for vancomycin, 0.064 μg.mL ^-1^ for cefepime, and incubated for 1 hour. Following incubation, the bacterial culture was centrifuged (6000x g, 5 minutes) and the antibiotic containing media was removed. The bacterial pellet was resuspended in pre-warmed media and serially diluted 1:10 in pre-warmed C+Y in a 96 well plate. Growth was measured every 20 minutes for 24 hours using a BioTek Synergy H1 (Absorbance: 620), and lag phase time was calculated by Agilent Gen6 data analysis software.

#### Murine immune treatment

All mouse experiments were performed with approval of and in accordance with guidelines of the St. Jude Institutional Animal Care and Use Committee. To simulate different immune states in our mouse model, female BALB/c mice from Jackson Laboratory were treated prior to pneumococcal inoculation. For neutrophil-depleted mice, roughly 200 mg/kg of cyclophosphamide was administered via intraperitoneal injection 120 hours prior to infection and a second injection of roughly 150 mg/kg was administered 48 hours prior to infection. For macrophage-depleted lineages, clodronate liposomes were intranasally administered to mice in a dose of 12.5 mg/kg 48 hours prior to infection.

#### *in vivo* co-infection

To determine the competitive advantage *in vivo*, wildtype TIGR4 and each mutant strain were grown in C+Y to logarithmic phase OD_620_ ∼0.6 – 0.8 and frozen as glycerol stocks at –80°C. Prior to infection, the glycerol stocks were thawed, washed, resuspended in 1× Phosphate buffered saline, and enumerated to confirm the correct CFUs infection dose. Seven-week-old female BALB/c mice were anesthetized with 2.5% isoflurane and intranasally inoculated with 100µL containing 10^6^ CFU of a 1:1 ratio of TIGR4 and pneumococcal mutant. For RNase Y competition experiments, mice were administered either vancomycin, cefepime, or levofloxacin via intraperitoneal injection 16 hours post-infection. Mice were euthanized either 22 hours post-infection for the RNase Y competition experiment or 24 hours post-infection for the nicotinamidase competition experiment using 3 L/min CO_2_ followed by cervical dislocation. Whole lungs and chest cavity blood were subsequently collected via dissection. Bacterial burden was enumerated by serially diluting into 1× PBS and plating either blood or homogenized lung supernatant onto tryptic soy blood agar plates, and bacterial titers were compared via Mann-Whitney U test.

#### Experimental Evolution *in vivo* Passaging

Bacterial populations were evolved *in vivo* in the presence of various antibiotics from the ancestral TIGR4 strain. BALB/c mice from Jackson Laboratories were intranasally challenged with 100 µL containing approximately 10^6^ CFU of TIGR4. Eight hours post infection, mice were subsequently treated with a single intraperitoneal injection of antibiotic. Mice were euthanized and lungs were removed via dissection, washed briefly with PBS and resuspended in 1 mL of PBS. Lung tissue was homogenized and rough tissue was pelleted via centrifugation at 300x g for 5 minutes. Bacteria-containing lung supernatant was then titered onto 20µg/mL neomycin, 3% blood agar plates for pneumococcal CFU counting. Lung titers were averaged by treatment and plotted with standard deviation at each passage. The remaining supernatant was plated onto additional plates for bacterial outgrowth after overnight incubation at 37°C. The overnight growth was harvested from plates into 15mL of C+Y media and made into 1mL aliquot frozen stocks in 25% glycerol. Roughly 10mL of the remaining bacterial cultures was pelleted via centrifugation (6000x g, 10 minutes) for use in genomic DNA preparation. Frozen aliquots were titered for CFU and used for the subsequent next generation of *in vivo* passaging. The antibiotic doses used to treat infected mice were either increased periodically or kept steady for selective pressure. Antibiotic doses were either increased after every third passage (meropenem, rifampicin, penicillin, vancomycin) for 15 passages, held constant for 15 passages (imipenem, ciprofloxacin, azithromycin), or held steady for the first 15 passages and then increased after every third passages for the next 15 passages (cefepime, levofloxacin, linezolid).

#### Minimum Inhibitory Concentration

Evolved populations were tested for MIC of the antibiotic in which they were propagated using E-test strips (Liofilchem). Three replicate tests were performed per lineage and duplicate readings were averaged for each replicate. Fold change in MIC relative to the ancestor was calculated by dividing the MIC of the evolved population by the MIC of the ancestral strain.

For scRNA studies, Minimum inhibitory concentration (MIC) of antibiotics was determined in THY using the microdilution method. In brief, a total of 1–5 × 10□ CFU of mid-exponential-phase bacteria were suspended in 100□μL of culture and combined with 100□μL of freshly prepared medium containing a single antibiotic to generate final concentration gradients of cefepime (0.008–0.8□μg.mL□¹), levofloxacin (0.1–2□μg.mL□¹), rifampicin (0.005–1□μg.mL□¹), and vancomycin (0.1–0.8□μg.mL□¹). Reactions were assembled in 96-well microplates, with each concentration tested in triplicate. Bacterial growth was continuously monitored at 37□°C for 16□h using a BioTek BioSpa Live Cell Analysis System (Agilent). The MIC was defined as the lowest antibiotic concentration that completely inhibited detectable growth under these conditions.

#### Genomic DNA Extraction

Genomic DNA from pneumococcal populations was extracted using a modified protocol using the Wizard Genomic DNA purification kit. Briefly, the bacterial populations harvested from the lung supernatant of passaged lineages were pelleted by centrifugation at 6000 ×*g* for 10 minutes, and supernatant removed and stored at -80 °C until genomic DNA extraction. Thawed bacterial pellets were lysed in 60 µL 10% deoxycholate and 60 µL 10% SDS. The suspension was incubated until clear (∼30 minutes) at 37°C, mixed with 600 µL of Nuclei Lysis Solution, and incubated at 80 °C for 5 minutes. Contents were cooled to room temperature, transferred to 2 mL microfuge tubes containing 3uL of RNase A solution (4 mg/ml), and incubated at 37 °C for 60 minutes. After cooling to room temperature, 250 µL of Protein Precipitation Solution was added, vortexed, and samples incubated on ice for 5 minutes. Samples were centrifuged at 16000 ×*g* for 5 minutes and supernatant transferred to a fresh tube containing 600 µL of isopropyl alcohol, mixed, and centrifuged (16000 ×*g* for 5 minutes) to precipitate and collect gDNA. Genomic DNA was washed twice with 70% ethanol, dried briefly, and resuspended with 200 µL of nuclease-free water. Genomic DNA was stored at -80 °C for genome sequencing.

#### Whole genome sequencing and analysis

All extracted genomic DNA samples were prepped and sequenced in the Hartwell Center at St. Jude. Sequence libraries were barcoded according to the Nextera XT DNA Library Preparation kit via liquid handling robots and run on the Illumina HiSeq2000 platform as previously described ^58^. Sequences were trimmed using Trimmomatic version 0.36 with the following parameters: ILLUMINACLIP: 2:30:10 LEADING:3 TRAILING:3 SLIDINGWINDOW:4:15 MINLEN:36 ^59^. For variant calling analysis, we used the *S. pneumoniae* TIGR4 reference genome from GenBank (NC_003028.3). Read mapping and variant calling were performed using Breseq version 0.35.0 in polymorphism mode to call variants at intermediate allele frequencies^60^. To reduce false positive variant calls, we utilized a conservative frequency cutoff of 10%. In addition, we required three reads from each strand to support each variant using the --polymorphism-minimum-variant-coverage-each-strand and --consensus-minimum-variant-coverage-each-strand flags.

Following breseq analysis, we excluded samples with poor sequencing quality, including samples with less than 30X average read depth and samples with high numbers of mutations (>100,000). We also removed variants which were fixed in the ancestral strain, as these variants do not reflect mutations selected during our experimental conditions. Mutations which occurred at exclusively intermediate frequencies and conflicted with other mutation trajectories, and thus are likely mapping errors due to repetitive genome regions, were also manually removed (**Supplementary File 1**). Finally, to identify mutations which most contributed to fitness *in vivo*, we removed mutations whose frequencies did not sum to 100% over all timepoints within a lineage for those passaged 15 times, and 200% for lineages passaged 30 times. Although breseq reports evidence for new junctions within the reference genome that may reflect large structural variants and regions of missing coverage, these variants were not considered for this analysis. As such, large structural genomic rearrangements and acquisition of new genetic material are not considered within this study.

#### Genome sequencing analysis and visualization

All data analysis and visualization of whole genome sequencing data was completed using R version 4.0.5 with packages tidyverse, matrixStats, vegan, viridis, cowplot, ggrepel, and randomcoloR^61-68^. Scripts are available from https://github.com/michellescribner/spn-evo.

#### Human neutrophil purification

Human donor leukopaks were obtained from the Houston Blood Center. Equal volumes of blood and 0.9% NaCL, 3% Dextran (17-20mL each) were gently mixed and incubated at room temperature for 25 minutes. The top layer containing peripheral blood mononuclear cells and polymorphonuclear cells was transferred to a new 50 mL conical tube and centrifuged for 10 minutes at 500 ×g. Supernatant was removed and white blood cell pellets were resuspended in 30 mL of PBS and re-centrifuged (10 minutes, 500 ×g). Supernatant was again discarded and pellets were resuspended in 17 mL of Hank’s Balanced Salt Solution and filtered through a 70 µm nylon cell strainer. 17 mL of each strained resuspended cell mixture was gently overlayed onto 12.5 mL of Ficoll and centrifuged at room temperature without centrifugal braking (30 minutes, 500 ×g, 0 deceleration). Remove PBMC and Ficoll layers via pipetting and resuspend PMN pellet with 10 mL PBS and transfer to 20 mL additional PBS in a new 50 mL conical tube. Centrifuge (10 minutes, 500 ×g) and discard supernatant. Lyse remaining RBCs by adding and resuspending pellet in 9 mL ACK Lysis Buffer, incubating at room temperature for 2.5 minutes and quenching reaction with the addition of 1 mL PBS. Centrifuge again (10 minutes, 350 ×g), discard supernatant, and resuspend the pellet in RPMI+10mM HEPES+0.1% human serum albumin. Enumerate the PMNs via hemocytometer.

#### Lactate dehydrogenase release assay

To measure neutrophil death in the presence of either TIGR4 or the nicotinamidase pneumococcal mutant, fresh neutrophils isolated from human donors were co-incubated with each respective pneumococcal strain for 3 hours and LDH release measured using the CytoTox-ONE™ Homogenous Membrane Integrity Assay (Promega). Briefly, pneumococcal strains were grown to roughly OD_600_: ∼0.4, concentrated and resuspended in RPMI + 10mM HEPES + 0.1% human serum albumin so that 10 µL media contains roughly 2.5 x 10^6^ CFU (2.5 x 10^8^ CFU/mL). In 96-well plates, 2.5 x 10^5^ neutrophils in 90 µL of RPMI + 10mM HEPES + 0.1% human serum albumin added to each well. At timepoint 0, 10 µL of bacteria in the aforementioned concentration was added to each co-incubation well, to a multiplicity of infection of 10. Positive and negative control wells contained only neutrophils and media in equivalent 100 µL volumes as the sample wells. Negative controls neutrophil-only wells were measured at the end of study. Positive controls neutrophil-only wells were immediately treated with 1 µL of lysis buffer to benchmark LDH release at 100% lysis. Wells were co-incubated for the respective timepoints at 37 °C and 5% CO_2_ with a final timepoint at 3 hours. At each timepoint, plates were centrifuged at 1500 rpm for 5 minutes and 50 µL of supernatant was removed from respective sample wells and transferred to a black 96-well clear-bottom fluorescence assay plate. Additionally, the remaining sample volume was used for bacteria CFU titering and enumeration. For fluorescence reading signifying LDH release, 50 µL of Cyto-Tox ONE Reagent was added to all experiment wells, mixed briefly for 30 seconds by orbital shaking, and incubated for 10 minutes at room temperature, protected from light. To read fluorescence, 25 µL of Stop Solution was added to each well, shaken for an additional 10 seconds and read on an EnVision Nexus plate reader (PerkinElmer, Excitation: 560 nm, Emission: 590 nm)

#### Western Blot

TIGR4 and nicotinamidase mutant T4ΔSP_1583::PhSE B were grown at 37 °C and 5% CO_2_ in 10 mL of C+Y liquid media to an OD^600^ = 0.5 and adjusted to the same CFU, verified by serial dilution and colony titering (approximately 1 x 10^9^ CFU/mL). 1 mL of cells were pelleted by centrifugation (8000 ×g, 5 minutes) and supernatant was saved for sampling. Pellets were lysed with 50 µL each of 10% sodium deoxycholate and 10% SDS via incubation at 37 °C for 30 minutes. Whole cell lysates and supernatant of each sample was run on 4-12% NuPAGE™ Bis-Tris Gels at 80V for 2 hours and samples transferred to a nitrocellulose membrane via Power Blotter (Invitrogen). Transferred samples were incubated for 2 hours with 1:5000 dilutions of rabbit pneumolysin toxoid antibody (Tuomanen laboratory), washed with PBS+0.1% Tween-20, and overnight at 4 °C with anti-rabbit secondary antibody in 5% non-fat dried milk powder. Membranes were washed, developed using SuperSignal™ West Femto Maximum Sensitivity Substrate, and bands visualized on a ChemiDoc™ MP Imaging System.

#### Detection and quantification of NETs in murine lung

FFPE lung sections were stained for neutrophils as described in Albrengues *et al* ^69^. Briefly, sections were de-parrafinized using Histoclear (#HS-200, National Diagnostics) and rehydrated using a decreasing ethanol gradient. Slides underwent heat-induced epitope retrieval (HIER) in pre-warmed Tris-EDTA Buffer (10 mM Tris Base, 1 mM EDTA, 0.05% Tween 20, pH 9.0) for 8 minutes on “high” in a pressure cooker. Following HIER, slides were washed and blocked in Fc Receptor Blocker (#NB309, Innovex Biosciences) for 30 minutes at room temperature. Slides were then blocked and permeabilized in 1X block (0.1% Triton-PBS; 2.5% BSA, 5% donkey serum) for 1 hour at room temperature. Slides were then incubated with goat-anti-myeloperoxidase (#AF3667, R&D Systems) 1:100 and rabbit-anti-citrullinated histone 3 (#ab5103, Abcam) 1:250 in 0.5X block overnight at 4 °C. The following morning slides were washed with PBS and incubated with donkey anti-goat AlexaFluor488 (#A11055, Invitrogen) 1:150, donkey anti-rabbit AlexaFluor568 (#A10042, Invitrogen) 1:400 DAPI 1:500 diluted in 0.5X block for 1 hour at room temperature in the dark. Slides were then washed in PBS and mounted with coverslips using Invitrogen ProLong Diamond Antifade Mountant (#P36961, Invitrogen). Entire lungs were imaged using the tilescan function on a Leica DMi8 Thunder Imager inverted fluorescent microscope.

Images were computationally cleared using Leica instant computational clearing prior to NET and neutrophil quantification. Image pixels were classified and segmented using Ilastik ^70^. Pixels were classified and segmented into “NET” or “background”, “Neutrophil” or “background”, and “Nuclei” or “background” using distinct pixel classification files. NETs were identified in training images as triple colocalization of MPO, citrullinated histone H3, and DAPI as has been previously described ^71^. Neutrophils were identified in training images as colocalization of MPO and DAPI. No images from the training set were used for the analysis. Binary segmented imaged were quantified in ImageJ using “Analyze Particles” function with a 10 pixels2 threshold for NETs and 90 pixels2 threshold for neutrophils. Percent NET area was calculated by ((Total NET Area/Total Neutrophil Area) x 100) for each lung.

#### Temporal collection for bulk RNA-seq experiments

Starter cultures of T4 strains were inoculated into THY medium supplemented with 5 µL.mL^-1^ oxyrase and incubated at 37 °C with 5% CO□ until mid-exponential phase (OD_600_ of 0.4–0.5), corresponding to approximately two bacterial doublings. Cultures were then washed and resuspended in fresh THY medium at an OD_600_ of 0.1. Three biological replicates per strain were prepared under each condition: absence or presence of cefepime (0.064 µg.mL^-1^) or rifampicin (0.175 µg.mL^-1^), followed by incubation at 37 °C with 5% CO_2_. After four hours, OD_600_ and total culture volume were recorded. Cells were harvested by centrifugation at 4,000 rpm for 7 minutes at room temperature and washed once with phosphate buffered saline (1x PBS; Fisher Scientific, Waltham, MA), and pelleted again under the same conditions. Pellets were resuspended in fresh THY medium at the original OD_600_, and cultures were incubated under the same conditions to monitor recovery (Figure 1A).

For RNA isolation, 10–40 mL of culture were collected by centrifugation at 4,000 rpm at 4°C at specified time points: baseline (T0), 30, 60, 120 and 240 minutes during antibiotic exposure (−exp), and 120 and 240 minutes post-antibiotic treatment during recovery (−rec). Cell pellets were snap-frozen in liquid nitrogen and stored at −80 °C until extraction using the RNeasy Mini Kit (Qiagen). RNA integrity number (RIN) was assessed using the Agilent 4150 TapeStation System with RNA and RNA High Sensitivity ScreenTape reagents (Agilent, Santa Clara, CA), and RNA concentration was quantified by Qubit RNA High Sensitivity Assay (ThermoFisher, Waltham, MA). Additionally, colony-forming units (CFU.mL^-1^) were enumerated by serial dilution and plating on blood agar plates, while total cell counts were determined via microscopy using an INCYTO C-Chip™ Disposable Hemacytometer (SKC) featuring a Neubauer Improved grid with 100 μm chamber depth.

#### Probe set design and amplification

Probes were designed as previously described ^72,73^. Using the reference genome NC_003028.3, probe sequences were constructed to include an mRNA-complementary region of approximately 50 bp, flanked at the 5′ end by a PCR handle of 18–23 bp and at the 3′ end by a 30 nt poly(dA) tract. Terminal regions of each probe contained sequences enabling circularization and HindIII restriction enzyme digestion. This design yielded 9,380 probes of ∼131 bp, collectively covering the entire *S. pneumoniae* strain T4 transcriptome (**Supplementary File 1**). Primary probes (Twist Bioscience, San Francisco, CA) were synthesized in the same orientation as the coding strand. Rolling circle amplification (RCA) mediated by phi29 DNA-polymerase was performed in three cycles, each reversing probe polarity; the final RCA products therefore correspond to sequences complementary to the target transcripts. RCA products underwent HindIII digestion using a primer-directed double-stranded region to facilitate enzyme cleavage. Single-stranded protoprobes were subsequently purified using the NucleoSpin Gel and PCR Clean-up Kit (Macherey–Nagel, Düren, Germany), substituting Buffer NTC for NTI to optimize recovery of single-stranded DNA. Oligonucleotides sequences listed in **Supplementary File 1**.

#### Single-Cell RNA sample collection, library preparation and sequencing

Time-killing assays were performed as described for the bulk experiments. T4 WT and *rny*L77S were cultured in the presence of cefepime (2xMIC, 0.064 µg.mL^-1^), and samples were collected at timepoints: T0, 60, 120 and 240 minutes during antibiotic exposure (−exp), and 120 and 240 minutes post-antibiotic treatment during recovery (−rec). Additional time-killing assays were run using only *rny*L77S in the presence of cefepime (2xMIC, 0.064 µg.mL^-1^), levofloxacin (2xMIC, 2□μg.mL□¹), vancomycin (2xMIC, 0.76□μg.mL□¹), osmotic pressure (300 mM NaCl), or acidic condition (pH 4.2), and samples were collected at T0 and 240 minutes of antibiotic exposure. Optical density (OD_600_) was calibrated to cell count, and sample volumes were adjusted to correspond to the cell number equivalent to 1 mL of culture at an OD_600_ of 1. For cultures with lower density (e.g., OD_600_ of 0.5), proportionally larger volumes (2 mL) were collected to maintain consistent cell numbers across samples. Cell cultures were fixed using a 30 min incubation in 1% paraformaldehyde (final concentration) in a rotoflex tube rotator/mixer at 10 rpm at 10 °C (Thomas Scientific, Swedesboro, NJ). Formaldehyde-fixed samples were washed with 0.02% saline sodium citrate (SSC, Invitrogen) by gentle centrifugation (6,000*g*) for 3 min. After the wash, cell pellets were resuspended in 1 mL MAAM (4:1 V:V dilution of methanol to glacial acetic acid). Samples were kept at −20 °C for up to 2 days before further processing. For in situ probe hybridization, ∼50 μL of fixed sample (∼1.5 × 10^7^ cells) were centrifuged and washed once in 1x PBS (Fisher Scientific, Waltham, MA) to remove methanol and acetic acid. From this point on, PBS-S was used (1x PBS supplemented with 0.01 U.μL^-1^ SUPERase In RNase Inhibitor; SUPERase-IN, Thermo Scientific, Waltham, MA). After the wash step, cells were incubated in 200 μL PBS-S with 350 u μl^−1^ of CPL1 (Recombinant *Streptococcus* phage Cp-1, Biomatik; Ontario, Canada) for 30 min at room temperature. CPL1 concentration (20µL of ∼413 µg.mL^-1^ of CPL1 + 25µL TEL-RI + 1µl SUPERASE-In; TEL-RI, as previously suggested by Blattman & colleagues ^14^; Briefly, 100 mM Tris pH 8.0, 50 mM EDTA, 0.1 U/μL SUPERase In RNase Inhibitor) was optimized as previously suggested ^15^. After 30 min, cells were collected and washed with 500 μL PBS-tween (PBS with 0.1% Tween 20). Cells were then resuspended with 100 μL of probe binding buffer consisting of 5 × SSC, 30% formamide, 9 mM citric acid (pH 6.0), 0.1% Tween 20, 50 ug ml−1 heparin and 10% low molecular weight dextran sulfate (Molecular Instruments, Los Angeles, CA). Cell suspensions were placed in a 50 °C shaker-incubator and allowed to pre-equilibrate for 1 hour. After 1 hour, probes (previously heated to 96 °C for 2 minutes and let to cool down to 50 °C) were added at a final concentration of 80 ng.µL^-1^ (in a total probe volume of 40 µL) to each cell suspension and samples were left to incubate overnight with shaking at 300 rpm. After the overnight incubation samples were washed five times in prewarmed (50 °C) probe-wash solution (5 × SSC, 30% formamide, 9 mM citric acid pH 6.0, 0.1% Tween 20 and 50 µg ml-1 heparin; Molecular Instruments, Los Angeles, CA). Before encapsulation on the 10X device, cells were washed three times and diluted in 1x PBS-S plus 0.1% BSA (New England Biolabs, Ipswich, MA) to a final suspension ratio of 1,000 cells. µL^-1^ as suggested in the 10X Chromium instruction manual.

For microfluidic encapsulation and droplet generation reaction, Single-cell partitioning, barcoding and cDNA library generation was achieved using the 10X Genomics Chromium Controller with the ChromiumTM Next GEM Single Cell 3’ Kit v3.1 (10x Genomics, Pleasanton, CA). For GEM generation (10X microfluidic encapsulation), a master mix containing the following reagents was prepared: 19 μL of 4X ddPCR Multiplex Supermix (Bio-Rad, San Diego, CA), 3 μL of custom primer (10X V3 indrop pcr1&2 Primer, 10 μM; **Supplementary File 1**), 2 μL Reducing Agent B (10X Genomics) and 7.8 μL molecular biology grade water (water was supplemented with BSA to a final concentration of 0.1%, MGW-B; Cornig, Glendale, AZ). Cells were resuspended in the master mix, and the suspensions were loaded into a 10x Chromium Controller (10x Genomics) with a targeted cell recovery of 10,000 cells.

After microfluidic partitioning on the 10×□Chromium Controller (10×□Genomics), each reaction was visually assessed under a microscope to confirm successful generation of Gel Bead-in-Emulsions (GEMs). Samples were then transferred to fresh PCR tubes and cycled at the following conditions: 94 °C for 5 min, six cycles of 94°C for 30 s followed by 50 °C for 30 s then 65 °C for 30 s, held at 4 °C. After PCR one, the emulsion was broken and the pooled DNA purified using NucleoSpin Gel and PCR Clean-up (Macherey–Nagel, Düren, Germany) as per manufacturers’ instructions for dsDNA. Purified DNA was amplified once more using a master mix composed of 17 μL of purified DNA, 20 μL of Q5 Hot Start 2X MM, 1.5 μL of forward primer (10x pcr2 fwd, 10 μM; **Supplementary File 1**) and 1.5 μL of 10X V3 indrop pcr1&2 (10 μM). PCR conditions included a 30 s incubation at 98 °C followed by 16 cycles of 10 s at 98 °C, 20 s at 62 °C and 20 s at 72 °C before a final extension at 72 °C for 2 min. After PCR two, amplified DNA was purified using the NucleoSpin Gel and PCR Clean-up Kit (Macherey-Nagel). Purified DNA was run on Agilent TapeStation to confirm the presence of a band at ∼200 bp, indicating successful library generation. Libraries were then prepared for sequencing by Illumina adaptor addition via low cycle (*n* = 6) PCR with custom library preparation finishing primers (**Supplementary File 1**). PCR 3 libraries were purified using a Select-a-Size DNA Clean & Concentrator Kit (Zymo Research, Irvine, CA), to obtain a single ∼230bp product per sample library. Sequencing was carried out on an AVITI 2x150 Sequencing Kit Cloudbreak FS High Output (Element Bioscience, San Diego, CA) that allows to obtain 1 billion reads per run. Final libraries were spiked with 2% PhiX and sequenced for 8 bp in each index, 28 bp Read 1 and 90 bp Read 2.

#### Processing of scRNA sequencing data

Sequencing reads were processed using the 10x Genomics Cell Ranger pipeline (v8.0.1). A custom reference was built from probe annotation files. Reads were aligned and counted with cellranger count using the custom reference. Spurious UMIs were removed as described previously ^72^, and normalization factors were estimated from the number of spurious UMIs. For each gene, the maximal probe count was used as the gene-level count. The top 10,000 cells with the highest total counts were retained for downstream analysis.

#### scRNA clustering and visualization

All single-cell analyses were performed in Scanpy (version 1.11.4) ^74^. Gene expression matrices were normalized to a total of 1 × 10□ counts per cell and log-transformed (base 10). Highly variable genes were identified using the cell ranger method (top 2,000 genes) and used for downstream analysis. Expression values were scaled to a maximum of 10 before performing principal component analysis (PCA). A neighborhood graph was constructed using 40 principal components and 10 nearest neighbors. Two-dimensional embeddings were generated using UMAP (v0.5.9.post2) ^75^, and cell clusters were identified with the Louvain algorithm (v0.8.2) ^76^.

Cell abundance for each cluster was estimated from bulk colony-forming unit (CFU) counts at each time point, scaled by the relative abundance of that cluster in the single-cell data. Doubling time was calculated from the fold change in estimated cell abundance between consecutive time points, normalized by the actual time interval.

#### RNA dynamics analysis

To estimate RNA abundance, gene expression matrices were normalized to a sequencing depth of one million reads per sample. The RNA content of each cell was then calculated as the sum of its normalized counts.

Normalized degradation and recovery rates were calculated in two steps. First, the sample-wise degradation or recovery rate was defined based on the median of total RNA content per cell in each sample, with the ratio of median RNA content between samples representing the sample-wise rate. Secondly, the Average transcript abundance per gene was computed as the mean normalized expression level across the top 10,000 cells per sample. Gene-wise degradation or recovery rates were defined as the ratio of mean transcript abundance between samples, normalized by the corresponding sample-wise rate.

Statistical significance of differences in degradation or recovery rates between the *rny*L77S and wild-type samples, or between defined gene groups, was assessed using a two-sided Mann–Whitney U test.

#### Entropy calculation

To quantify transcriptional variability, we calculated entropy based on the variance of log□ fold changes (log□FCs) relative to corresponding NDC samples. As the majority of NDC cells were assigned to cluster 3, these cluster 3 NDC cells were used as the reference. Treated sample subgroups were then compared to this reference to assess differential expression using a Wilcoxon rank-sum test. Genes with p-values < 0.05 were included in the entropy calculation. Entropy was defined following the previous study ^35^:

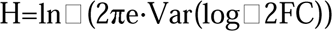

where Var(log□2FC) is the variance of the log□ fold changes.

#### Temporal trajectory inference

Dynamic cell-state transitions in the *rny*L77S strain were modeled using TemporalProblem framework in moscot (v0.4.3; ^77^, with cells annotated by time post-treatment. A RealTimeKernel from CellRank (v2.0.7; ^36^ was constructed to estimate transition probabilities between cell states, and transition matrices were computed considering all self-transitions, with edge weights scaled and thresholded to retain biologically meaningful connections.

To explore potential transitions between states over time, starting populations of cells were selected from each Louvain-defined cluster at initial time points. Random walk simulations were performed iteratively for multiple stop times, with 10,000 simulations per stop time and a maximum of 500 iterations. Transition probabilities were visualized on the force-directed layout, providing a map of likely cell-state transitions in the mutant population throughout the experiment.

#### Inter-cluster distance analysis

To assess divergence and convergence of mutant cell populations over time, pairwise Euclidean distances were computed between all cells from clusters 2 and 3 at each time point using their UMAP embeddings.

#### Gene set enrichment analysis

To identify biological pathways that exhibited significantly different degradation or recovery rates between the *rny*L77S mutant and the wild type, preranked gene set enrichment analysis (GSEA) was performed using the GSEApy Python package (v1.1.10; ^78^). For each pair of time points, per-gene degradation or recovery rate differences were computed as the rate in *rny*L77S minus that in the corresponding wild-type sample. GSEA was conducted against Gene Ontology biological process gene sets (annotation information can be found in “Annotation_TIGR4.tsv” in Mendeley Data), with a minimum and maximum gene set size of 10 and 200, respectively. Statistical significance was determined based on 1,000 permutations of the ranked gene list.

## SUPPLEMENTAL INFORMATION, TITLES, AND LEGENDS

**Figure S1.** A) Dose titration of various antibiotics against pneumococcal bacterial burdens in murine lungs 24 hours post-infection. Incremental doses of antibiotics were administered to mice after intranasal infection of mice. Antibiotic doses were increased until an approximate one log drop in recoverable lung bacterial CFUs was observed. B) *S. pneumoniae* populations evolved in mice differing in immunocompetence with antibiotic selection. Lung titers (CFU/lung homogenate) were measured at every passage. Mean and standard deviation for the three parallel lineages per immune treatment are shown. Antibiotic concentrations used throughout the evolution experiment are shown beneath each drug. C) Counts of types of unique mutations within all evolved populations. Each distinct mutation is counted only once. D) Number of mutations acquired by each bacterial lineage that passed filtering parameters, including those which were outcompeted prior to the final passage. E) Instances of parallel evolved genes in which five or more mutations evolved independently across populations. F) Number of samples with specific *rny* (RNase Y) mutations by antibiotic treatment.

**Figure S2. Jaccard similarity matrix of genetic diversity among evolved lineages.** (A) Higher coefficient indicates greater degree of overall genetic similarity in each respective lineage. Limited similarity was observed among lineages grouped either by host immune state or by drug mechanism of action. The exception to this observation is rifampicin-evolved passages, which are more similar because they share on-target resistance mutations, in *rpoB*. VNC = vancomycin, PEN = penicillin, MEM = meropenem, LNZ = linezolid, LVX = levofloxacin, IPM = imipenem, CIP = ciprofloxacin, CEF = cefepime, AZM = azithromycin, IPM=imipenem, RIF=rifampicin, M0 = macrophage-depleted mouse immune state, N0 = neutrophil-depleted mouse immune state, Nd = non-depleted (wildtype) mouse immune state.

**Figure S3. Allele frequency dynamics within evolved populations from all ten antibiotic regimens.** The frequencies of mutations within key genes that arose in parallel and reached high frequencies are shown at each mouse passage. A single passage represents the recovered pneumococcal population harvested from the lungs of a single mouse after infection and treatment with a respective antibiotic. Az = azithromycin, Ce = cefepime, Ci = ciprofloxacin, Im = imipenem, Le = levofloxacin, Li = linezolid, Me = meropenem, Pe = penicillin, Ri = rifampicin, Va = vancomycin, M0 = macrophage-depleted mouse immune state, N0 = neutrophil-depleted mouse immune state, Nd = non-depleted (wildtype) mouse immune state.

**Figure S4. *in vivo* and *in vitro* phenotypes of the nicotinamidase pneumococcal mutant.** (A) Pneumococcal CFU were enumerated 24 hours post-infection from the blood of mice intranasally challenged with a 1:1 ratio of the nicotinamidase mutant T4Δ1583::PhSE and the wildtype TIGR4 strain. For each mouse the competitive index was calculated as the CFU/mL of the mutant divided by the CFU/mL of the wildtype. Indices greater than one indicate a competitive advantage for the mutant strain. (B) LDH release of human neutrophils as a measure of neutrophil death during incubation with either TIGR4 or the nicotinamidase mutant. C) Pneumococcal growth and CFU concentrations in neutrophil LDH assay. During co-incubation with human donor neutrophils, cultures were sampled, diluted and plated at each respective timepoint to enumerate either TIGR4 or nicotinamidase mutant colonies present in the growth medium. D) Western blot visualization of pneumolysin present in TIGR4 versus nicotinamidase mutant. Cultures of either TIGR4 or the nicotinamidase mutant were grown to an OD_600_ of 0.4 and titered to ensure similar CFU counts. Whole cell lysates and supernatants of cultures were primary labeled with a 1:2000 dilution of pneumolysin toxoid L460D antisera from rabbit. E) Immunohistochemistry image quantitation of total and apoptotic neutrophils present in the lungs of infected mice. Murine lungs were excised from euthanized mice 24 hours post-infection and stained for cleaved caspase 3 to visualized apoptotic neutrophils in the lung. Pneumococcal CFU in the lungs were also enumerated at this time. F) Immunohistochemistry image quantification of neutrophil extracellular traps in the lungs of infected mice. Murine lungs were excised from euthanized mice 24 hours post-infection and stained for citrullinated histone H3 and myeloperoxidase to assess NET area relative to total neutrophils.

**Figure S5. Antibiotic tolerance screening of final passages of *in vivo* evolved pneumococcal populations.** Bacterial CFUs were quantified after 4 hours of exposure to either 1×, 2×, or 4× MIC of the final recovered pneumococcal population. Antibiotic killing activity is depicted as the log reduction in growth of the bacterial population compared to untreated population growth at each respective timepoint. Sample populations are shown in orange hue while the TIGR4 ancestral strain is shown in purple.

**Figure S6. Fold-change in MIC of RNase Y point mutants compared to parent TIGR4**. Mutant strain MICs were obtained for each antibiotic in which each respective mutation arose. Mutations occurring at identical locations but in separate antibiotic backgrounds were also tested for cross-resistance to the respective antibiotic. Striped bars indicate the originating antibiotic for each point mutant. Error bars represent the standard deviation of three replicate MICs.

**Figure S7. Antibiotic kill curves for RNase Y point mutants**. Graphs depict bacterial CFU/mL counts of RNase Y point mutants compared to the parent TIGR4 strain with and without levofloxacin (orange), cefepime (blue), and vancomycin (green). Antibiotic concentrations used were 2× the determined MIC for each respective strain. Colored lines indicate the evolved population while black lines indicate the original TIGR4 ancestral strain. Triangles represent the drug-exposed kill curves for evolved and ancestral strains. Squares and circles symbolize the untreated growth curves for evolved and ancestral strains, respectively.

**Figure S8. Known genomic diversity and structural clustering of alleles of the *S. pneumoniae rny* locus.** (A) Conservation of *rny* across the maximum likelihood phylogeny of 1,969 pneumococcal whole genomes obtained from the Monocle database (https://data-viewer.monocle.sanger.ac.uk/project/gps) compiled by the Global Pneumococcal Sequencing (GPS) project (https://www.pneumogen.net/gps/). The outer ring surrounding the phylogeny indicates the presence or absence of the RNase Y gene in pneumococcal genomes, confirming its conservation across the species. (B) Distribution of RNase Y protein alleles, i.e., unique versions of the amino acid sequences, across the whole-genome phylogeny of the pneumococcal genomes. For clarity, the outer ring surrounding the phylogeny is annotated by the ten most common alleles. (C) A chord diagram showing the correspondence of the RNase Y alleles and the genetic background of the pneumococcal strains. For clarity, only the four most common alleles are shown with everything else grouped as “other” alleles. (D) A chord diagram showing the correspondence of the RNase Y alleles and the susceptibility to the beta-lactam antibiotics of the pneumococcal strains inferred genotypically based on the genomic sequences. For clarity, only the four most common alleles are shown with everything else grouped as “other” alleles. (E) Mapping of the allelic variants from the evolution experiments (highlighted in red) and those observed in clinical isolates (highlighted in green) based on AlphaFold predictions.

**Figure S9. Single-cell transcriptomic data allow characterization of RNA content per cell and cell fate.** (A) Distribution of per-cell RNA content for WT cells in Clusters 1, 3, and 4 at each time point. Dashed lines indicate median values. *P*-values from two-sided Mann–Whitney U tests comparing Clusters 1 and 4 are shown in the plot. (B) Results of 10,000 random walk simulations based on the CellRank transition model constructed using the RealTimeKernel from 1h-exp to 2h-exp (top), 4h-exp (middle), and 4h-rec (bottom) initiated from Cluster 2 (left) or Cluster 3 (right) cells. Black dots denote starting points, and yellow dots indicate endpoints.

**Figure S10. RNA integrity number (RIN)**. RNA samples from Bulk experiments were adjusted to an optimal RNA functional range (25-500 ng.µL^-1^) and analyzed using RNA ScreenTape (TapeStation, Agilent). RNA integrity screentapes are shown in each of the three biological replicates (top, middle and bottom panels), for *S. pneumoniae* T4 WT (A) and *rny*L77S (B), in the absence (NDC) or presence of cefepime (CEF) or (RIF) during antibiotic exposure (-exp) and recovery (-rec). RIN values are high and kept at high levels in the *rny*L77S during the entire experiment (during and after antibiotic exposure), while WT RIN values decrease faster during the antibiotic treatment and do not increase during the recovery period.

**Figure S11. Enhanced decay and recovery of ribosomal transcripts drive a general tolerance response in *rny*L77S.** (A-B) Normalized degradation rates from T0 to 2h-exp plotted against log10 CPM in T0 for WT (A) and *rny*L77S (B). (C-D) Normalized degradation rates from T0 to 4h-exp plotted against log10 CPM in T0 for WT (C) and *rny*L77S (D). Rates were normalized to the sample-wise degradation rate in WT. (E-F) Normalized recovery rates from 4h-exp to 4h-rec plotted against log10 CPM in T0-NDC for WT (E) and *rny*L77S (F). Rates in A-F were normalized to the sample-wise degradation rate in WT. All Pearson correlation P values in the related plots are < 4.26×□10^□5^. (G-H) Gene set enrichment analysis (GSEA) of GO biological process gene sets based on ranked differences in normalized degradation or recovery rates between WT and *rny*L77S during exposure (G) and recovery (H). NES: Normalized enrichment scores.

**Fig S12. Convergence between clusters supports dual survival strategies and highlights the role of ribosomal transcripts in tolerance responses in *rny*L77S.** (A-B) Normalized degradation rates from T0 to 2h-exp plotted against log10 CPM in T0 for *rny*L77S Cluster 2 (A) and Cluster 3 (B). (C-D) Normalized degradation rates from T0 to 4h-exp plotted against log10 CPM in T0 for *rny*L77S Cluster 2 (C) and Cluster 3 (D). (E-F) Normalized recovery rates from 4h-exp to 4h-rec plotted against log10 CPM in T0 for *rny*L77S Cluster 2 (E) and Cluster 3 (F). Rates in A-F were normalized to the sample-wise degradation rate in *rny*L77S. (G-H) UMAP for 12 pooled samples showing Cluster 2 (G,I,K,M,O) and Cluster 3 (H,J,L,N,P) cells colored by SP_0210 (log_10_ CPM: G-H), SP_0213 (RNA content: I-J, log_10_ CPM: K-L), and SP_0214 (RNA content: M-N, log_10_ CPM: O-P). Blue oval highlights *rny*L77S cells from recovery samples; red oval highlights *rny*L77S cells from exposure samples.

**Figure S13. Single-cell transcriptomic data shows global RNA decay as a general tolerance response in *rny*L77S.** (A) UMAP visualization of six pooled single-cell transcriptomes from time zero no drug control (T0), 4h exposure to CEF, vancomycin (VNC), levofloxacin (LXV), high osmolarity (300 mM NaCl), and acidic conditions (pH 4.2) conditions, colored by sample. (B) Normalized degradation rates from T0 to 4h-exp were plotted against the log10 CPM in T0 for the cefepime (CEF) condition. Pearson correlation P-value = 0.

**Supplementary File 1.** Mutations manually curated due to biologically implausible allele frequency trajectories. RNA decay rates between TIGR4 and *rny*L77S. TIGR4 transcriptomic probe set. Single-stranded protoprobe sequences used for single-cell RNA-seq. Finishing primer sequences used in custom library preparation. Primers used for mutant construction.

