## Supplementary Material for "RNA Quality Control Enables Antibiotic Tolerance"

**Figure S1.** A) Dose titration of various antibiotics against pneumococcal bacterial burdens in murine lungs 24 hours post-infection. Incremental doses of antibiotics were administered to mice after intranasal infection of mice. Antibiotic doses were increased until an approximate one log drop in recoverable lung bacterial CFUs was observed. B) *S. pneumoniae* populations evolved in mice differing in immunocompetence with antibiotic selection. Lung titers (CFU/lung homogenate) were measured at every passage. Mean and standard deviation for the three parallel lineages per immune treatment are shown. Antibiotic concentrations used throughout the evolution experiment are shown beneath each drug. C) Counts of types of unique mutations within all evolved populations. Each distinct mutation is counted only once. D) Number of mutations acquired by each bacterial lineage that passed filtering parameters, including those which were outcompeted prior to the final passage. E) Instances of parallel evolved genes in which five or more mutations evolved independently across populations. F) Number of samples with specific *rny* (RNase Y) mutations by antibiotic treatment.

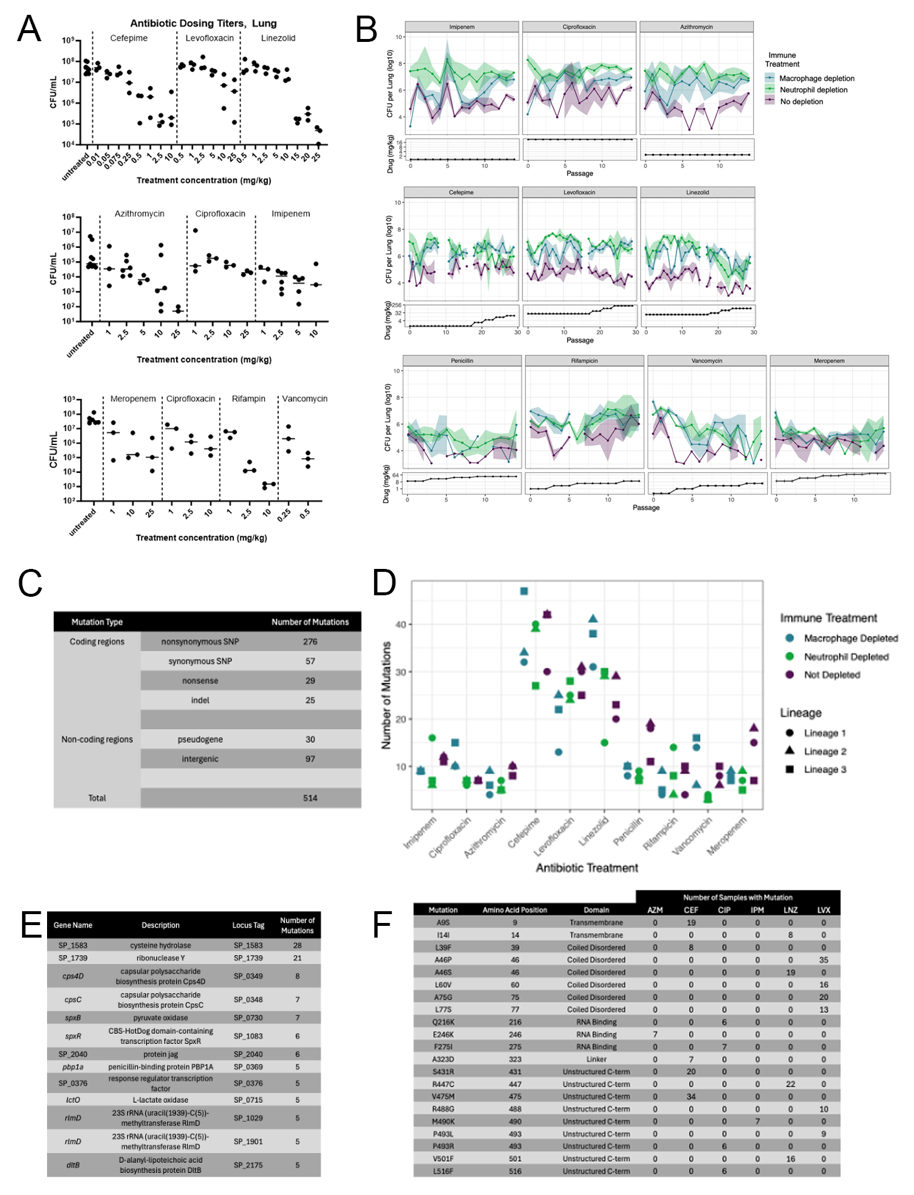

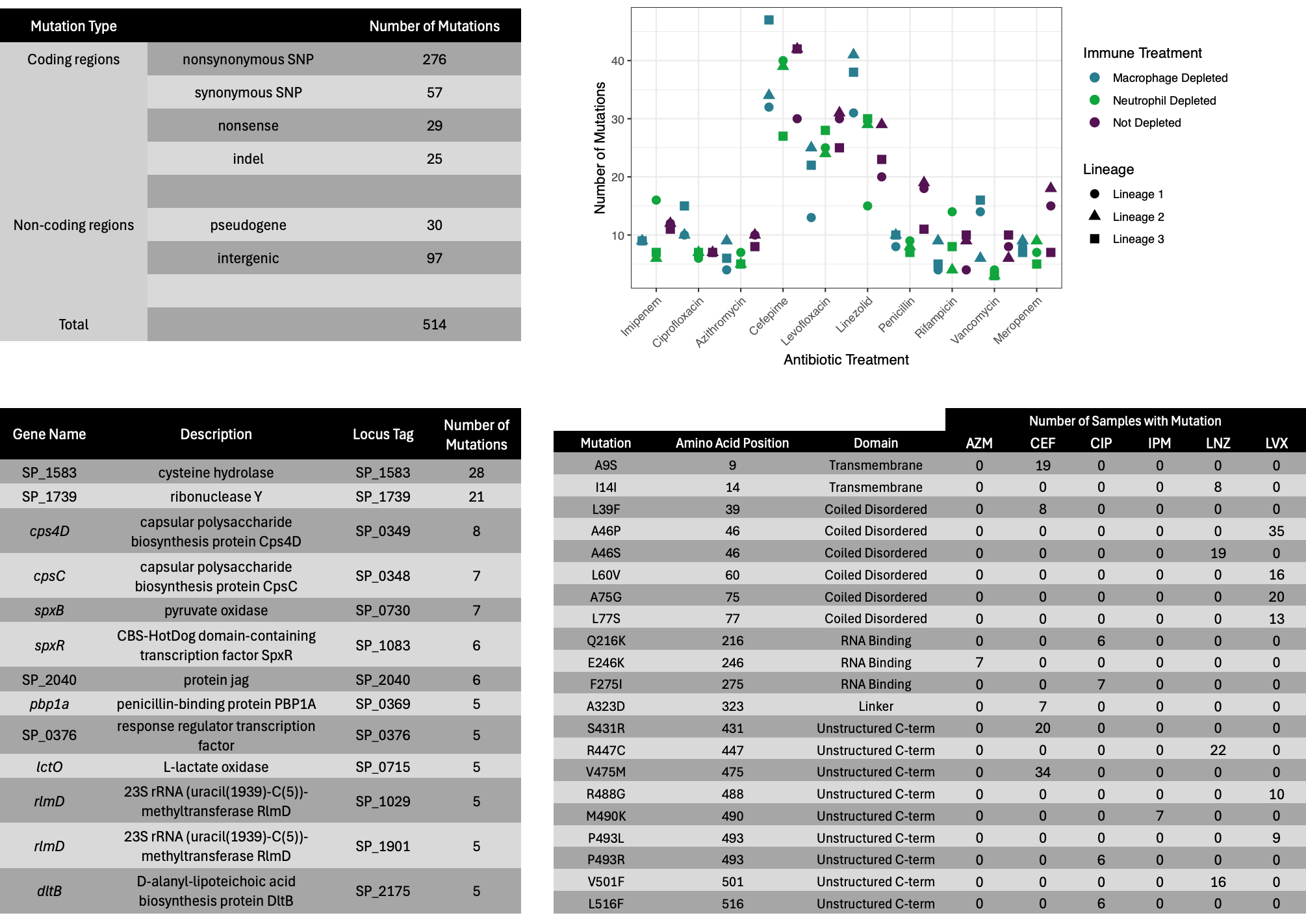

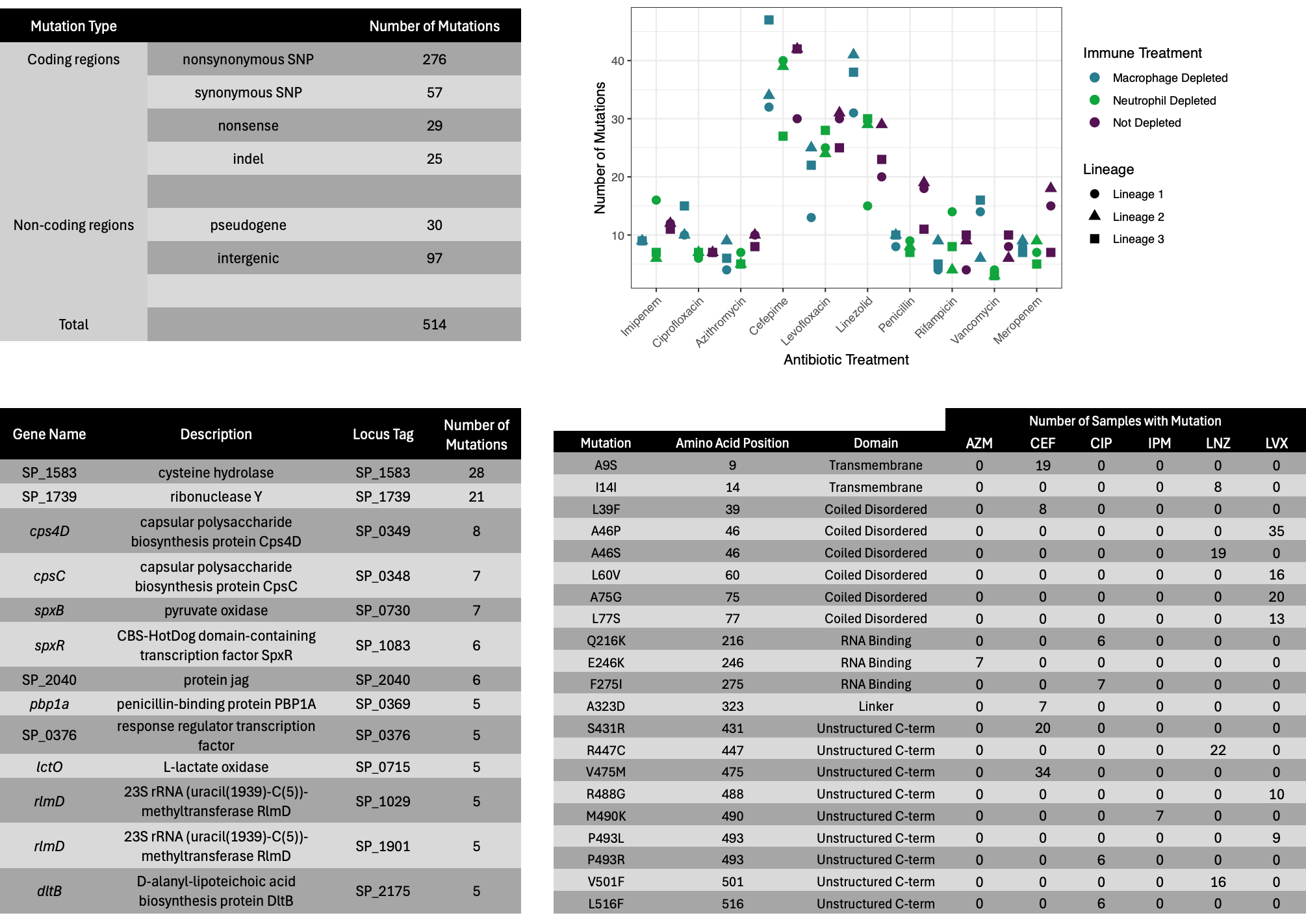

**Figure S2. Jaccard similarity matrix of genetic diversity among evolved lineages.** (A) Higher coefficient indicates greater degree of overall genetic similarity in each respective lineage. Limited similarity was observed among lineages grouped either by host immune state or by drug mechanism of action. The exception to this observation is rifampicin-evolved passages, which are more similar because they share on-target resistance mutations, in *rpoB*. VNC = vancomycin, PEN = penicillin, MEM = meropenem, LNZ = linezolid, LVX = levofloxacin, IPM = imipenem, CIP = ciprofloxacin, CEF = cefepime, AZM = azithromycin, IPM=imipenem, RIF=rifampicin, M0 = macrophage-depleted mouse immune state, N0 = neutrophil-depleted mouse immune state, Nd = non-depleted (wildtype) mouse immune state.

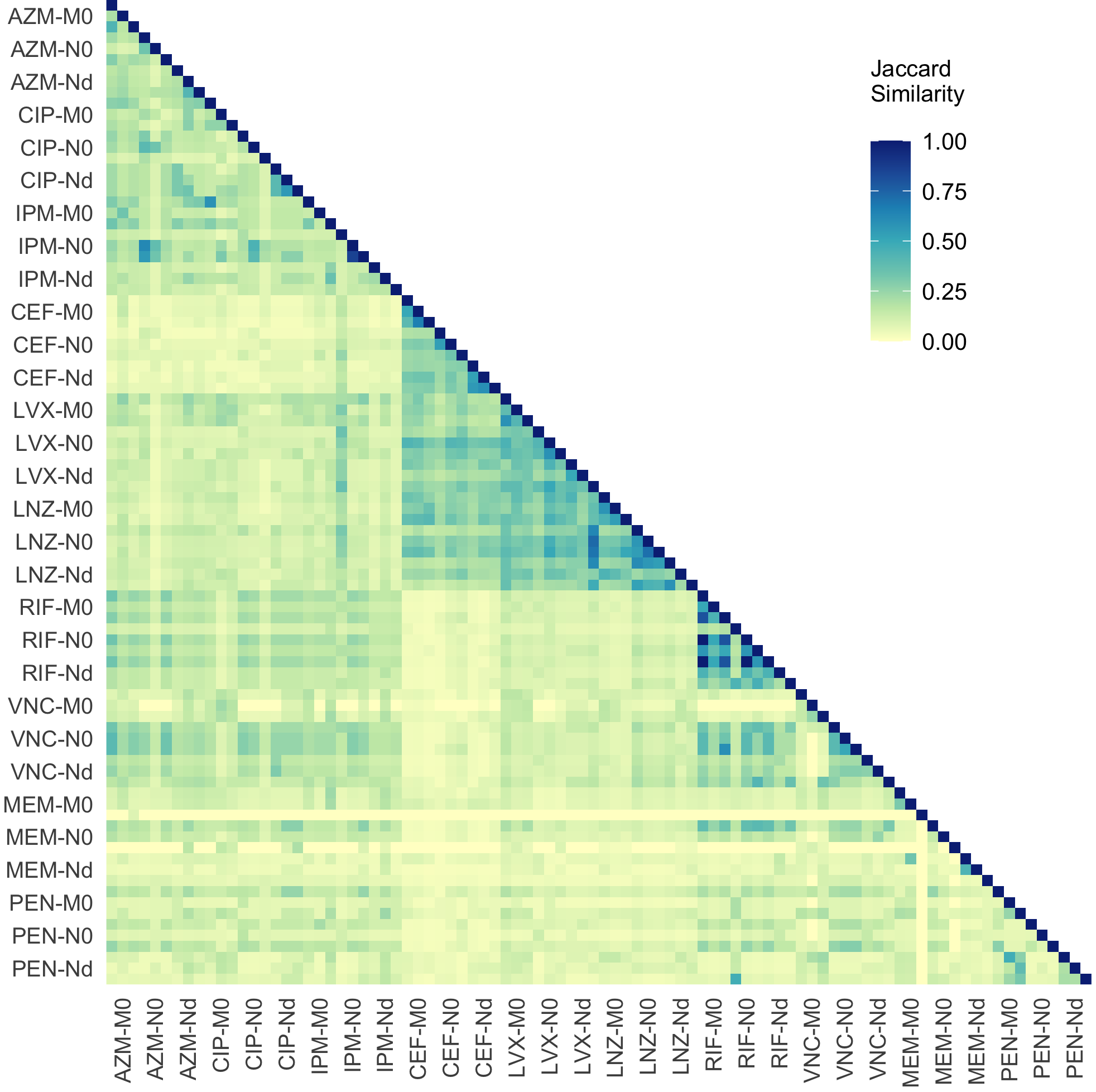

**Figure S3. Allele frequency dynamics within evolved populations from all ten antibiotic regimens.** The frequencies of mutations within key genes that arose in parallel and reached high frequencies are shown at each mouse passage. A single passage represents the recovered pneumococcal population harvested from the lungs of a single mouse after infection and treatment with a respective antibiotic. Az = azithromycin, Ce = cefepime, Ci = ciprofloxacin, Im = imipenem, Le = levofloxacin, Li = linezolid, Me = meropenem, Pe = penicillin, Ri = rifampicin, Va = vancomycin, M0 = macrophage-depleted mouse immune state, N0 = neutrophil-depleted mouse immune state, Nd = non-depleted (wildtype) mouse immune state.

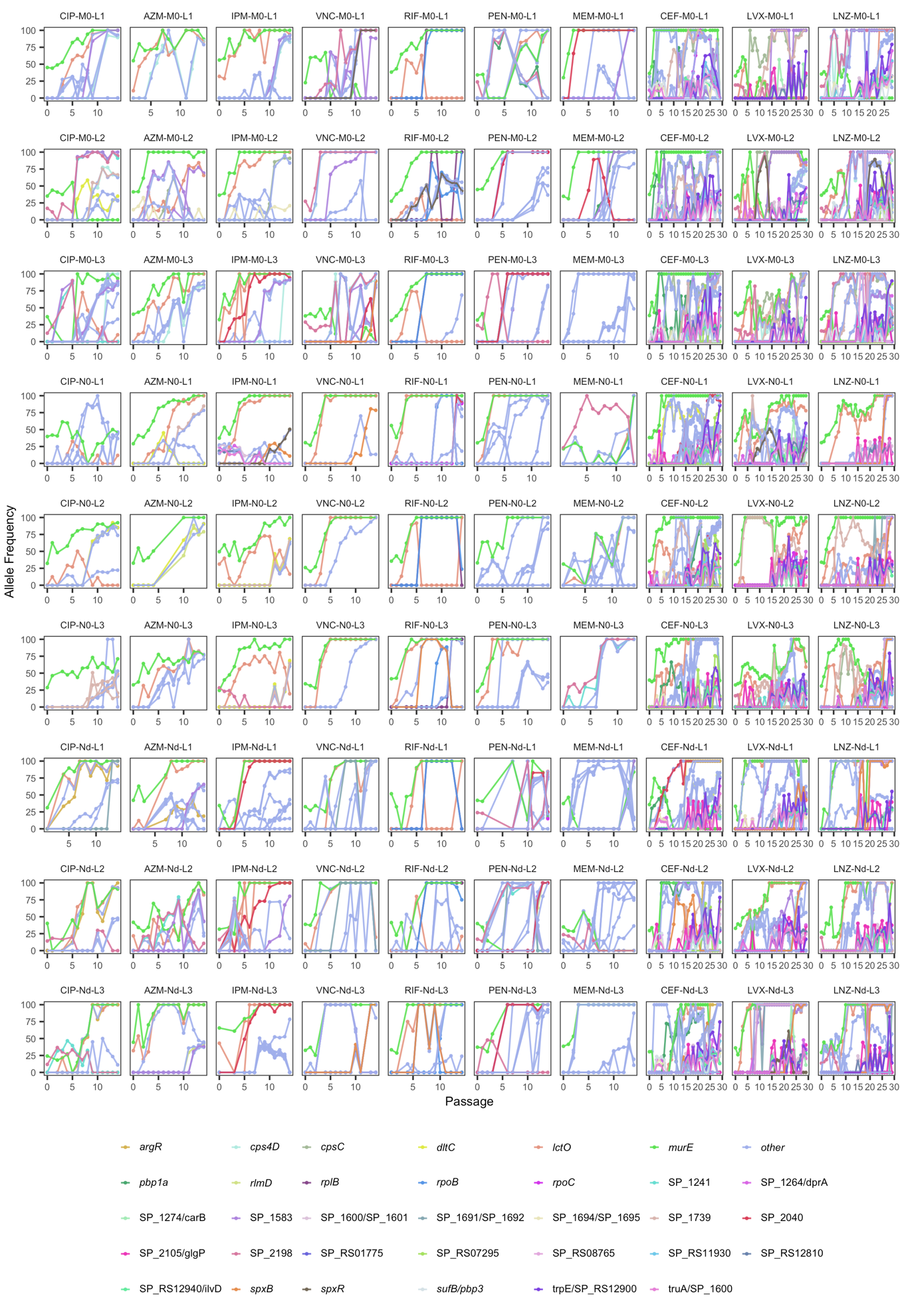

**Figure S4. *in vivo* and *in vitro* phenotypes of the nicotinamidase pneumococcal mutant.** (A) Pneumococcal CFU were enumerated 24 hours post-infection from the blood of mice intranasally challenged with a 1:1 ratio of the nicotinamidase mutant T4∆1583::PhSE and the wildtype TIGR4 strain. For each mouse the competitive index was calculated as the CFU/mL of the mutant divided by the CFU/mL of the wildtype. Indices greater than one indicate a competitive advantage for the mutant strain. (B) LDH release of human neutrophils as a measure of neutrophil death during incubation with either TIGR4 or the nicotinamidase mutant. C) Pneumococcal growth and CFU concentrations in neutrophil LDH assay. During co-incubation with human donor neutrophils, cultures were sampled, diluted and plated at each respective timepoint to enumerate either TIGR4 or nicotinamidase mutant colonies present in the growth medium. D) Western blot visualization of pneumolysin present in TIGR4 versus nicotinamidase mutant. Cultures of either TIGR4 or the nicotinamidase mutant were grown to an OD_600_ of 0.4 and titered to ensure similar CFU counts. Whole cell lysates and supernatants of cultures were primary labeled with a 1:2000 dilution of pneumolysin toxoid L460D antisera from rabbit. E) Immunohistochemistry image quantitation of total and apoptotic neutrophils present in the lungs of infected mice. Murine lungs were excised from euthanized mice 24 hours post-infection and stained for cleaved caspase 3 to visualized apoptotic neutrophils in the lung. Pneumococcal CFU in the lungs were also enumerated at this time. F) Immunohistochemistry image quantification of neutrophil extracellular traps in the lungs of infected mice. Murine lungs were excised from euthanized mice 24 hours post-infection and stained for citrullinated histone H3 and myeloperoxidase to assess NET area relative to total neutrophils.

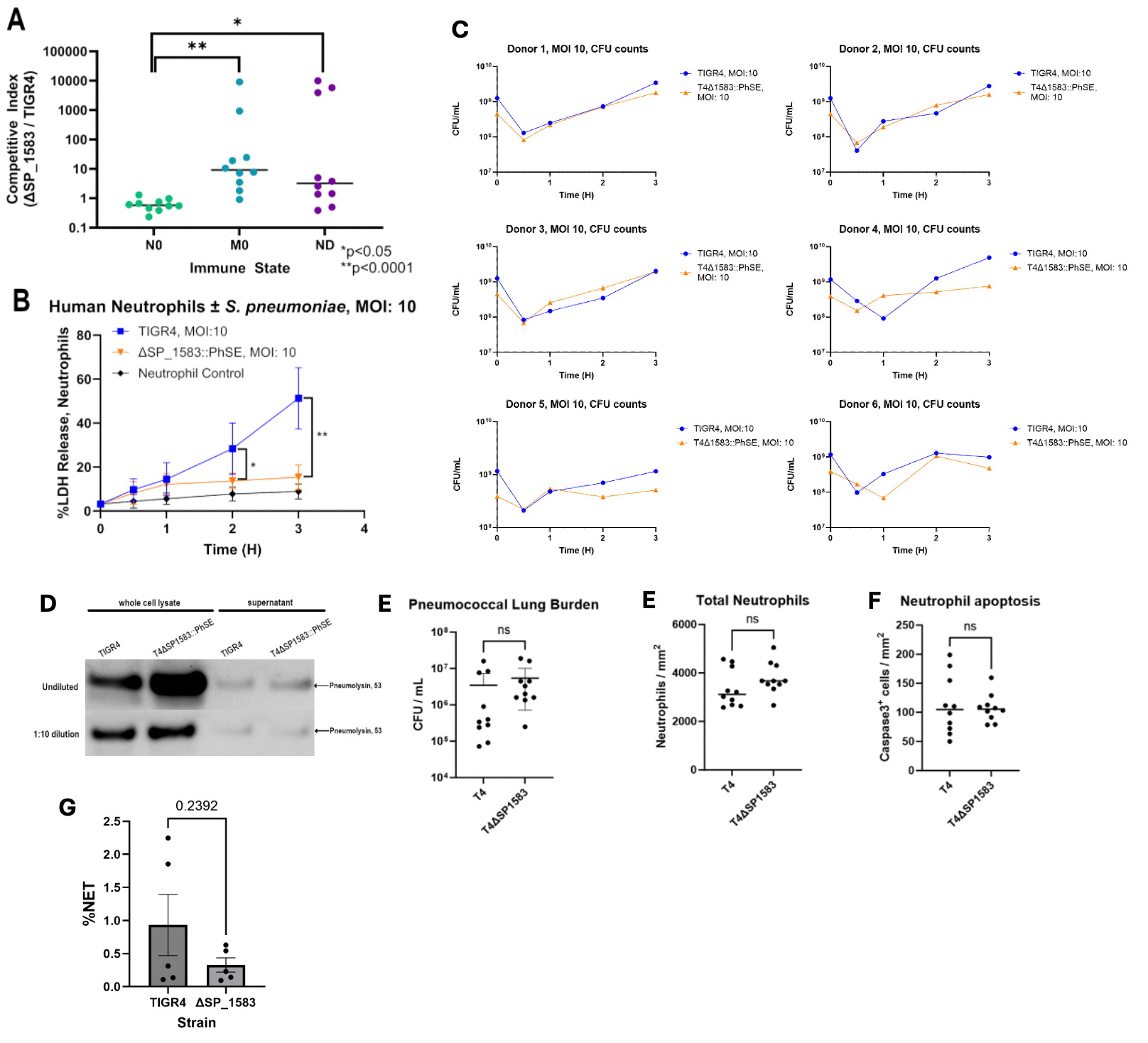

**Figure S5. Antibiotic tolerance screening of final passages of *in vivo* evolved pneumococcal populations.** Bacterial CFUs were quantified after 4 hours of exposure to either 1×, 2×, or 4× MIC of the final recovered pneumococcal population. Antibiotic killing activity is depicted as the log reduction in growth of the bacterial population compared to untreated population growth at each respective timepoint. Sample populations are shown in orange hue while the TIGR4 ancestral strain is shown in purple.

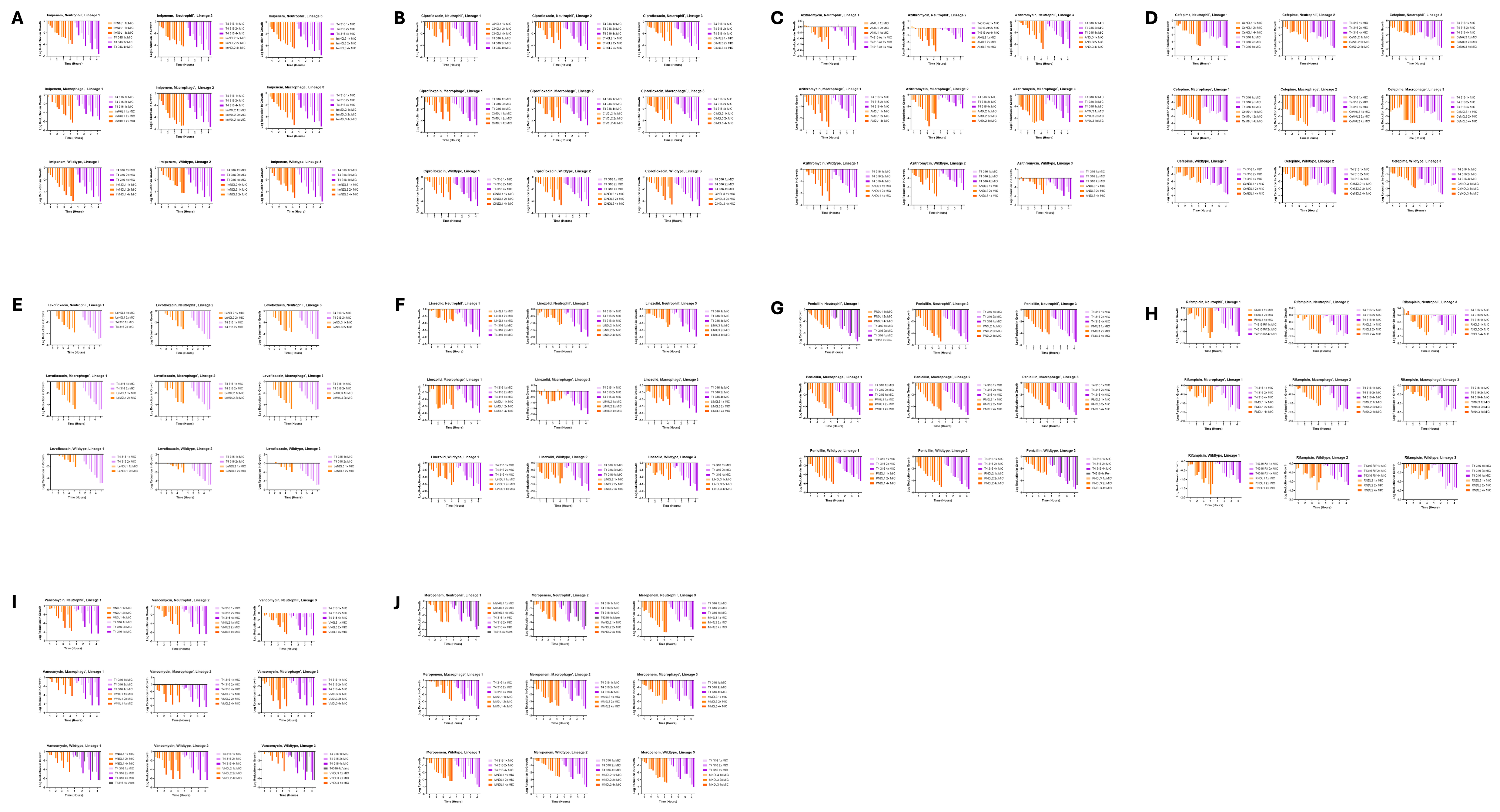

**Figure S6. Fold-change in MIC of RNase Y point mutants compared to parent TIGR4**. Mutant strain MICs were obtained for each antibiotic in which each respective mutation arose. Mutations occurring at identical locations but in separate antibiotic backgrounds were also tested for cross-resistance to the respective antibiotic. Striped bars indicate the originating antibiotic for each point mutant. Error bars represent the standard deviation of three replicate MICs.

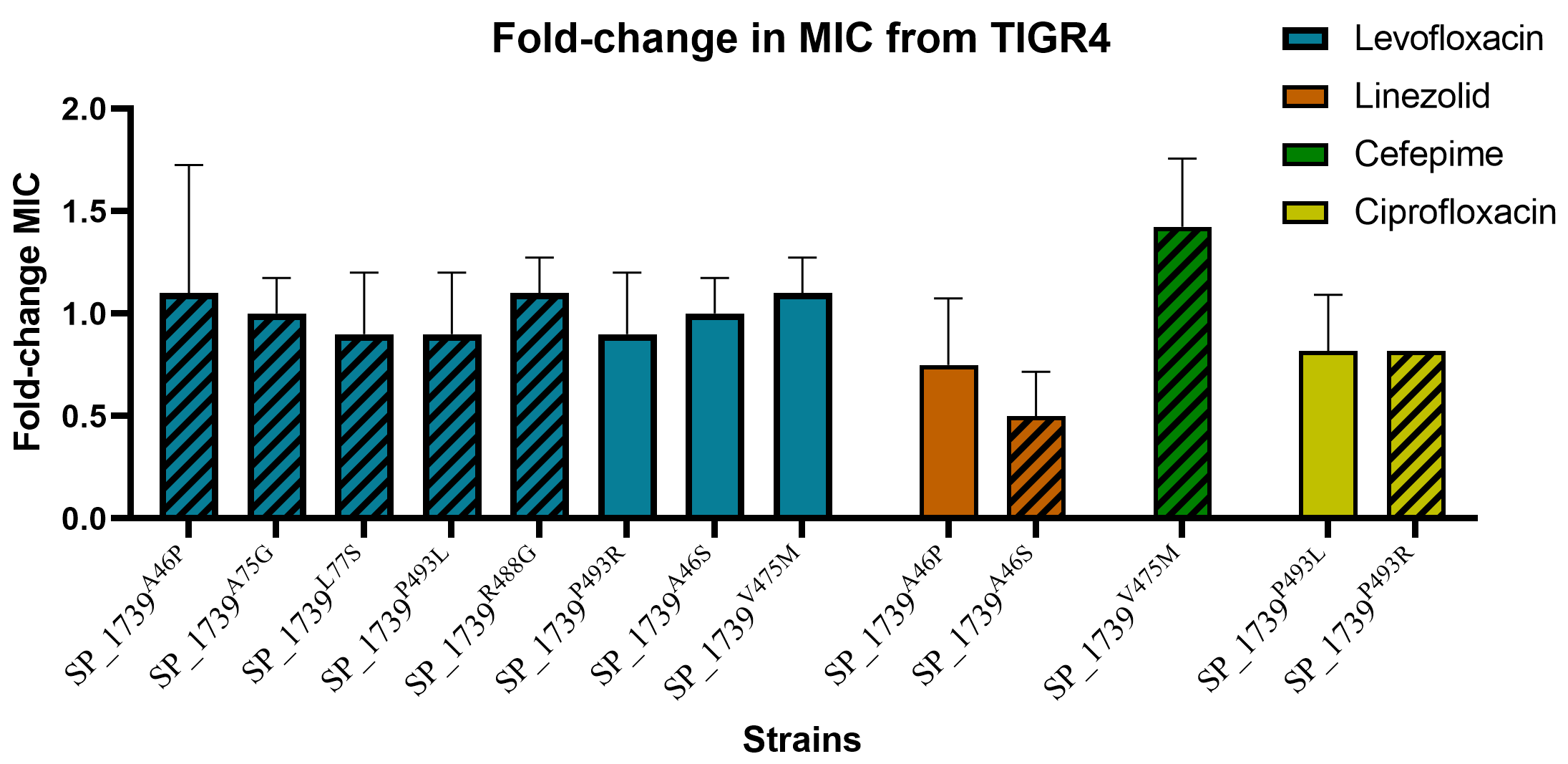

**Figure S7. Antibiotic kill curves for RNase Y point mutants**. Graphs depict bacterial CFU/mL counts of RNase Y point mutants compared to the parent TIGR4 strain with and without levofloxacin (orange), cefepime (blue), and vancomycin (green). Antibiotic concentrations used were 2× the determined MIC for each respective strain. Colored lines indicate the evolved population while black lines indicate the original TIGR4 ancestral strain. Triangles represent the drug-exposed kill curves for evolved and ancestral strains. Squares and circles symbolize the untreated growth curves for evolved and ancestral strains, respectively.

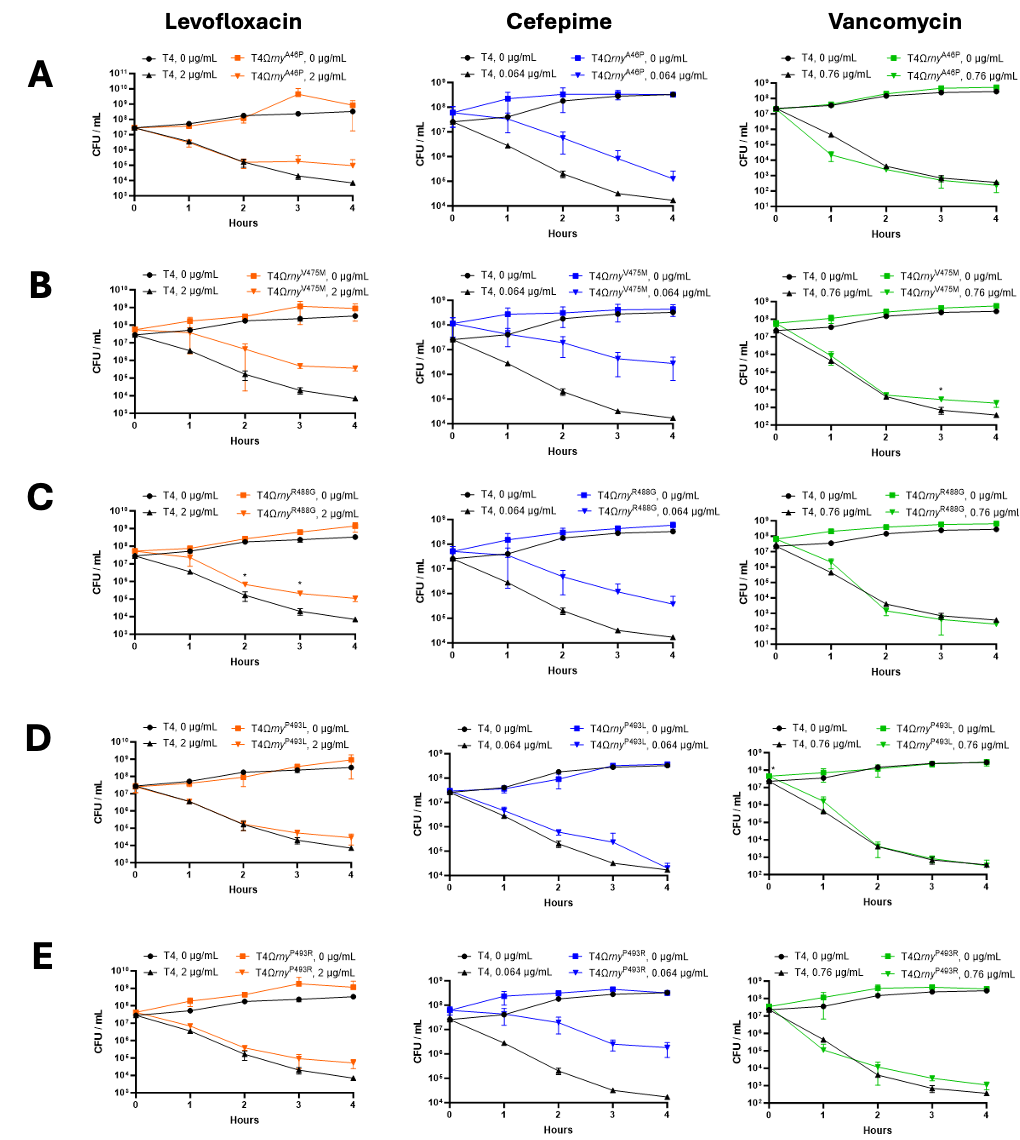

### **Figure S8.** **Known genomic diversity and structural clustering of alleles of the *S. pneumoniae* *rny* locus.** (A) Conservation of *rny* across the maximum likelihood phylogeny of 1,969 pneumococcal whole genomes obtained from the Monocle database (https://data-viewer.monocle.sanger.ac.uk/project/gps) compiled by the Global Pneumococcal Sequencing (GPS) project (https://www.pneumogen.net/gps/). The outer ring surrounding the phylogeny indicates the presence or absence of the RNase Y gene in pneumococcal genomes, confirming its conservation across the species. (B) Distribution of RNase Y protein alleles, i.e., unique versions of the amino acid sequences, across the whole-genome phylogeny of the pneumococcal genomes. For clarity, the outer ring surrounding the phylogeny is annotated by the ten most common alleles. (C) A chord diagram showing the correspondence of the RNase Y alleles and the genetic background of the pneumococcal strains. For clarity, only the four most common alleles are shown with everything else grouped as “other” alleles. (D) A chord diagram showing the correspondence of the RNase Y alleles and the susceptibility to the beta-lactam antibiotics of the pneumococcal strains inferred genotypically based on the genomic sequences. For clarity, only the four most common alleles are shown with everything else grouped as “other” alleles. (E) Mapping of the allelic variants from the evolution experiments (highlighted in red) and those observed in clinical isolates (highlighted in green) based on AlphaFold predictions.

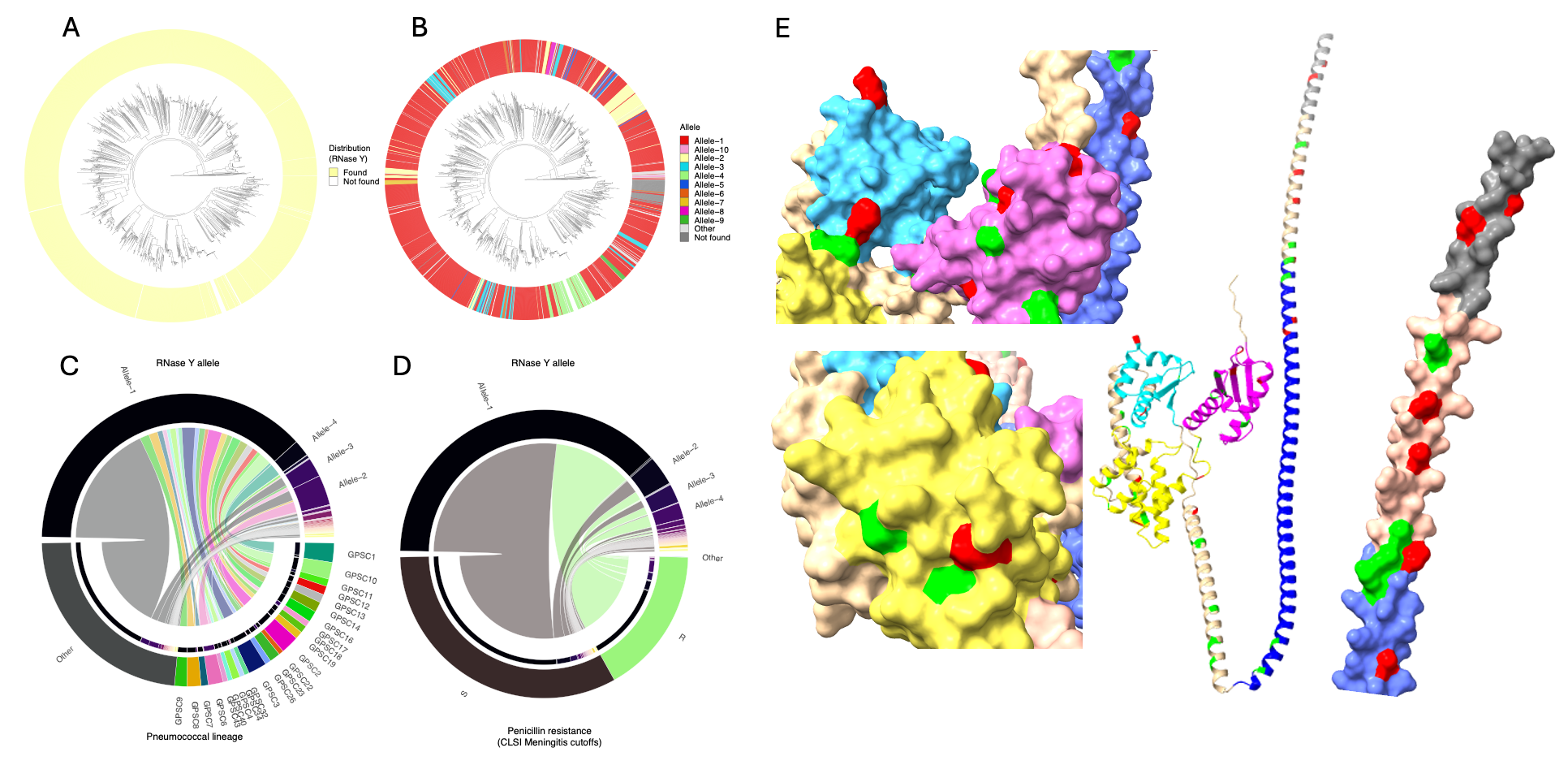

### **Figure S9. Single-cell transcriptomic data allow characterization of RNA content per cell and cell fate.** (A) Distribution of per-cell RNA content for WT cells in Clusters 1, 3, and 4 at each time point. Dashed lines indicate median values. *P*-values from two-sided Mann–Whitney U tests comparing Clusters 1 and 4 are shown in the plot. (B) Results of 10,000 random walk simulations based on the CellRank transition model constructed using the RealTimeKernel from 1h-exp to 2h-exp (top), 4h-exp (middle), and 4h-rec (bottom) initiated from Cluster 2 (left) or Cluster 3 (right) cells. Black dots denote starting points, and yellow dots indicate endpoints.

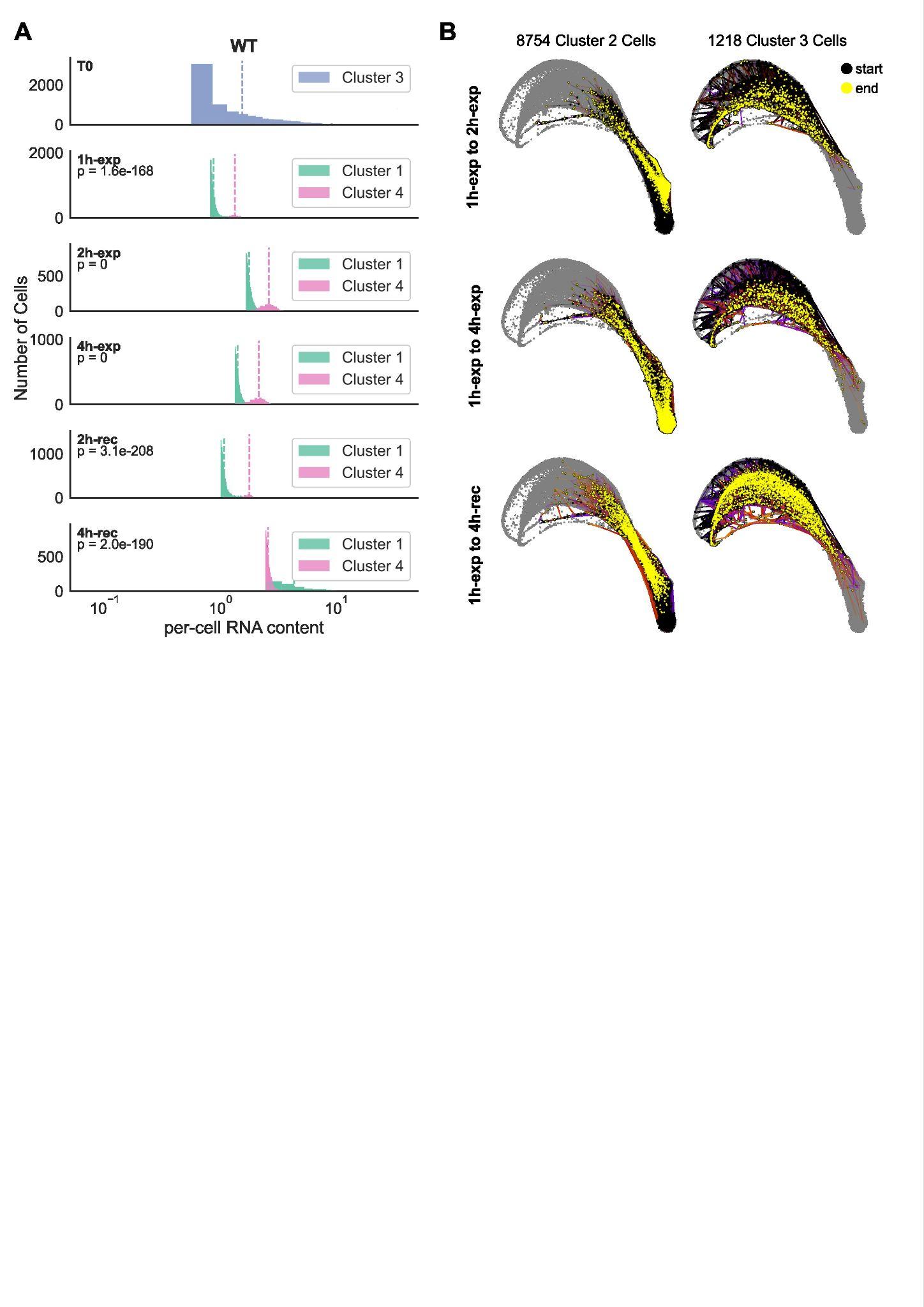

### **Figure S10. RNA integrity number (RIN)**. RNA samples from Bulk experiments were adjusted to an optimal RNA functional range (25-500 ng.µL^-1^) and analyzed using RNA ScreenTape (TapeStation, Agilent). RNA integrity screentapes are shown in each of the three biological replicates (top, middle and bottom panels), for *S. pneumoniae* T4 WT (A) and *rny*L77S (B), in the absence (NDC) or presence of cefepime (CEF) or (RIF) during antibiotic exposure (-exp) and recovery (-rec). RIN values are high and kept at high levels in the *rny*L77S during the entire experiment (during and after antibiotic exposure), while WT RIN values decrease faster during the antibiotic treatment and do not increase during the recovery period.

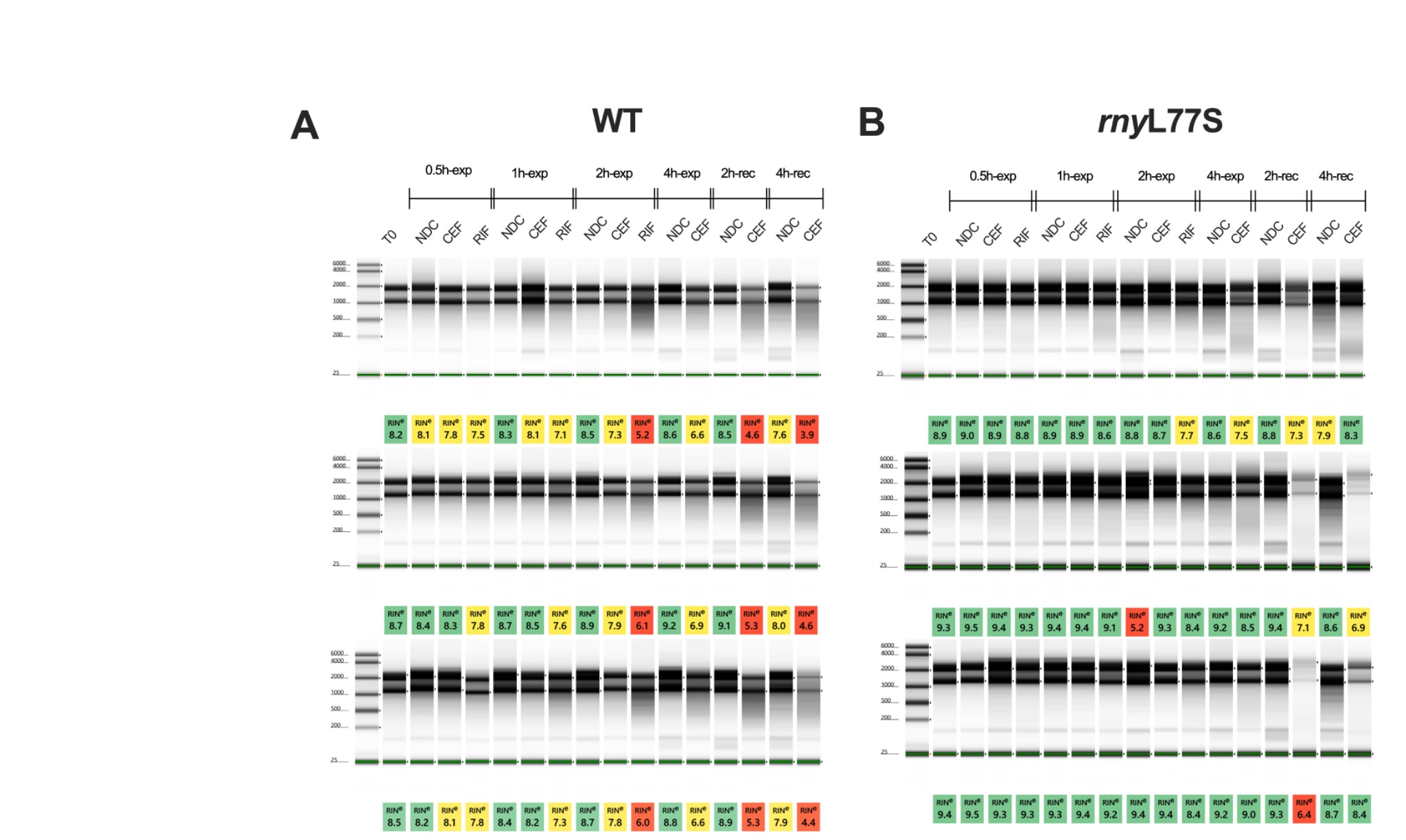

### **Figure S11. Enhanced decay and recovery of ribosomal transcripts drive a general tolerance response in *rny*L77S.** (A-B) Normalized degradation rates from T0 to 2h-exp plotted against log10 CPM in T0 for WT (A) and *rny*L77S (B). (C-D) Normalized degradation rates from T0 to 4h-exp plotted against log10 CPM in T0 for WT (C) and *rny*L77S (D). Rates were normalized to the sample-wise degradation rate in WT. (E-F) Normalized recovery rates from 4h-exp to 4h-rec plotted against log10 CPM in T0-NDC for WT (E) and *rny*L77S (F). Rates in A-F were normalized to the sample-wise degradation rate in WT. All Pearson correlation P values in the related plots are < 4.26× 10^⁻5^. (G-H) Gene set enrichment analysis (GSEA) of GO biological process gene sets based on ranked differences in normalized degradation or recovery rates between WT and *rny*L77S during exposure (G) and recovery (H). NES: Normalized enrichment scores.

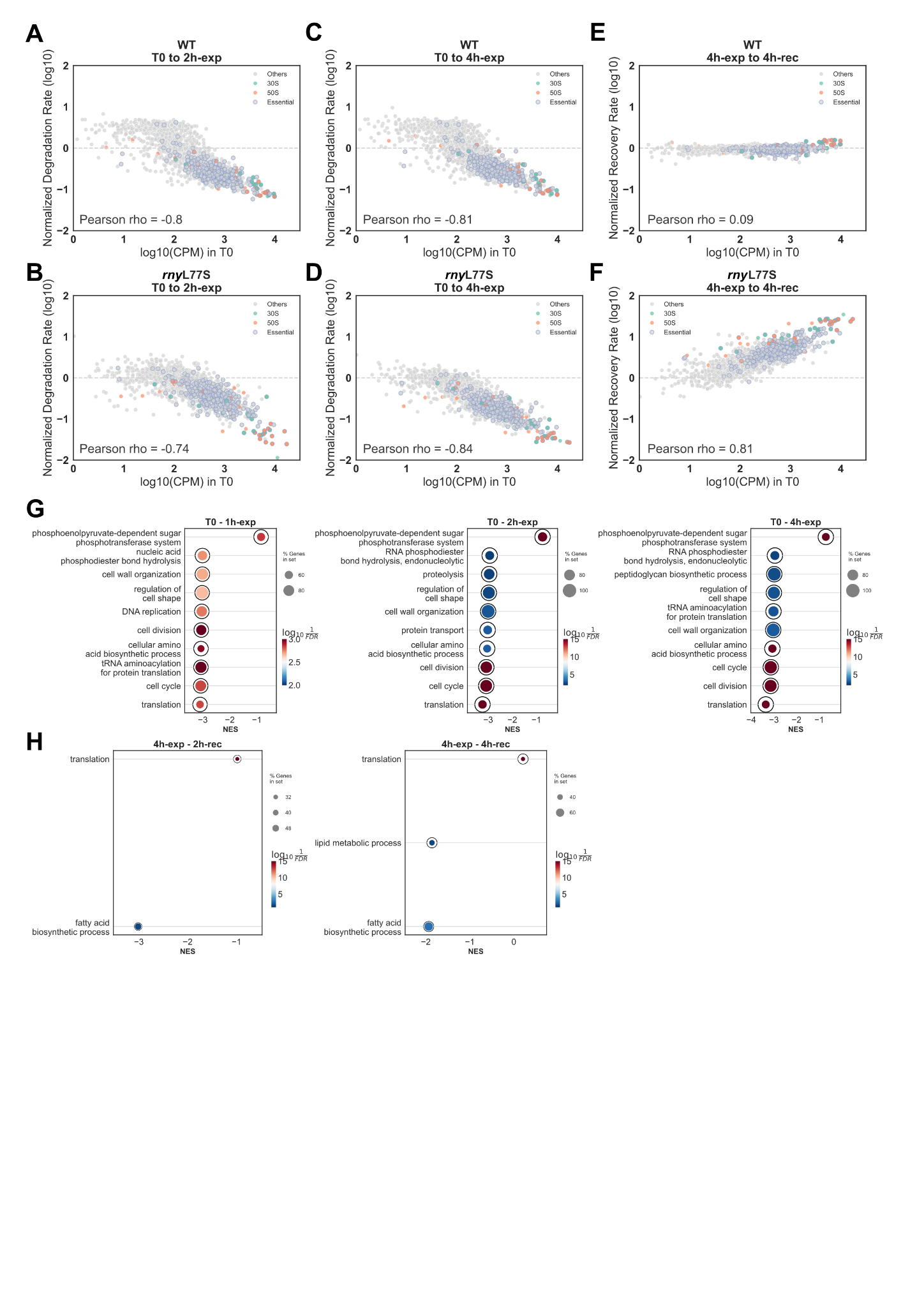

### **Fig S12. Convergence between clusters supports dual survival strategies and highlights the role of ribosomal transcripts in tolerance responses in *rny*L77S.** (A-B) Normalized degradation rates from T0 to 2h-exp plotted against log10 CPM in T0 for *rny*L77S Cluster 2 (A) and Cluster 3 (B). (C-D) Normalized degradation rates from T0 to 4h-exp plotted against log10 CPM in T0 for *rny*L77S Cluster 2 (C) and Cluster 3 (D). (E-F) Normalized recovery rates from 4h-exp to 4h-rec plotted against log10 CPM in T0 for *rny*L77S Cluster 2 (E) and Cluster 3 (F). Rates in A-F were normalized to the sample-wise degradation rate in *rny*L77S. (G-H) UMAP for 12 pooled samples showing Cluster 2 (G,I,K,M,O) and Cluster 3 (H,J,L,N,P) cells colored by SP_0210 (log_10_ CPM: G-H), SP_0213 (RNA content: I-J, log_10_ CPM: K-L), and SP_0214 (RNA content: M-N, log_10_ CPM: O-P). Blue oval highlights *rny*L77S cells from recovery samples; red oval highlights *rny*L77S cells from exposure samples.

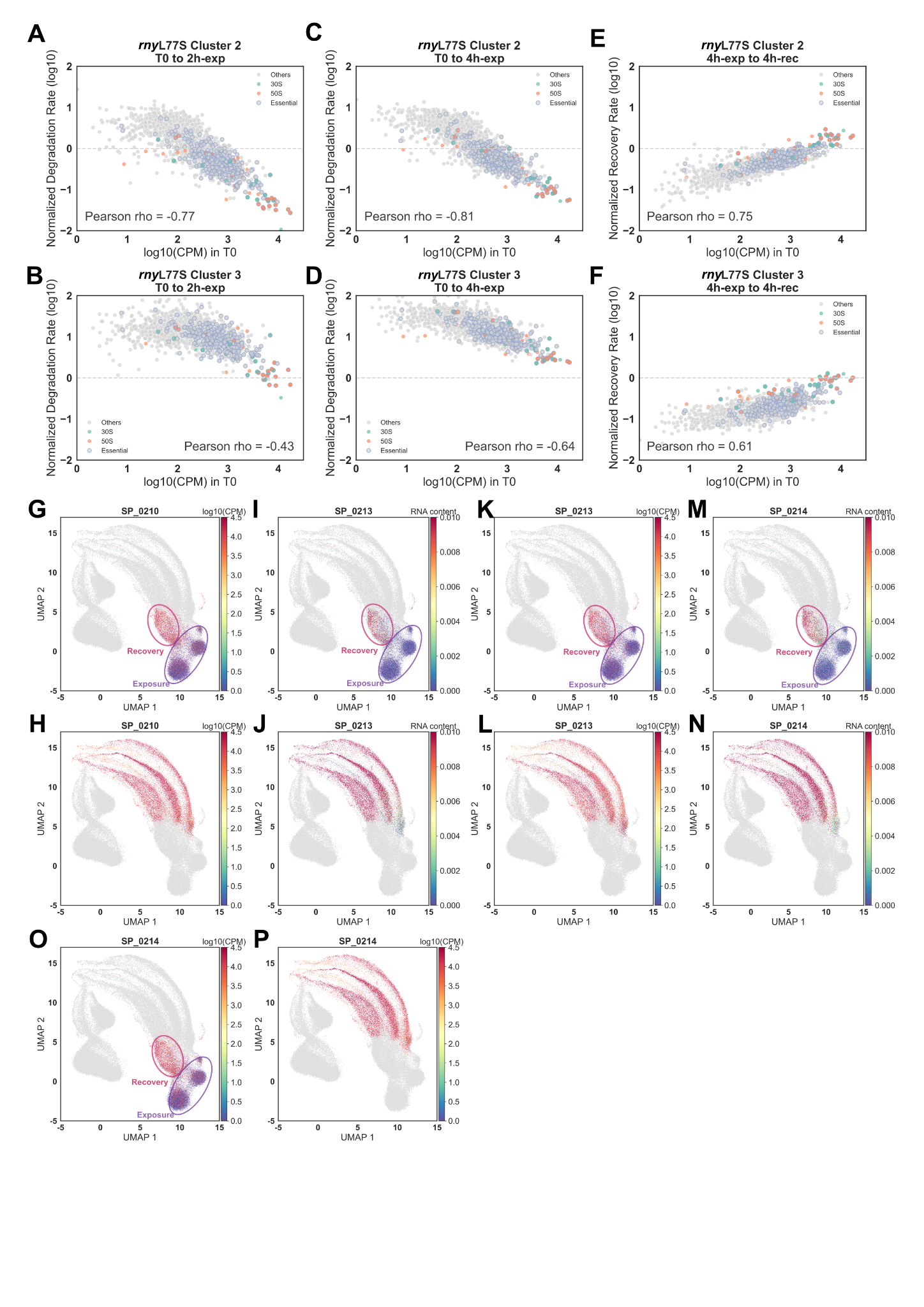

### **Figure S13. Single-cell transcriptomic data shows global RNA decay as a general tolerance response in *rny*L77S.** (A) UMAP visualization of six pooled single-cell transcriptomes from time zero no drug control (T0), 4h exposure to CEF, vancomycin (VNC), levofloxacin (LXV), high osmolarity (300 mM NaCl), and acidic conditions (pH 4.2) conditions, colored by sample. (B) Normalized degradation rates from T0 to 4h-exp were plotted against the log10 CPM in T0 for the cefepime (CEF) condition. Pearson correlation P-value = 0.

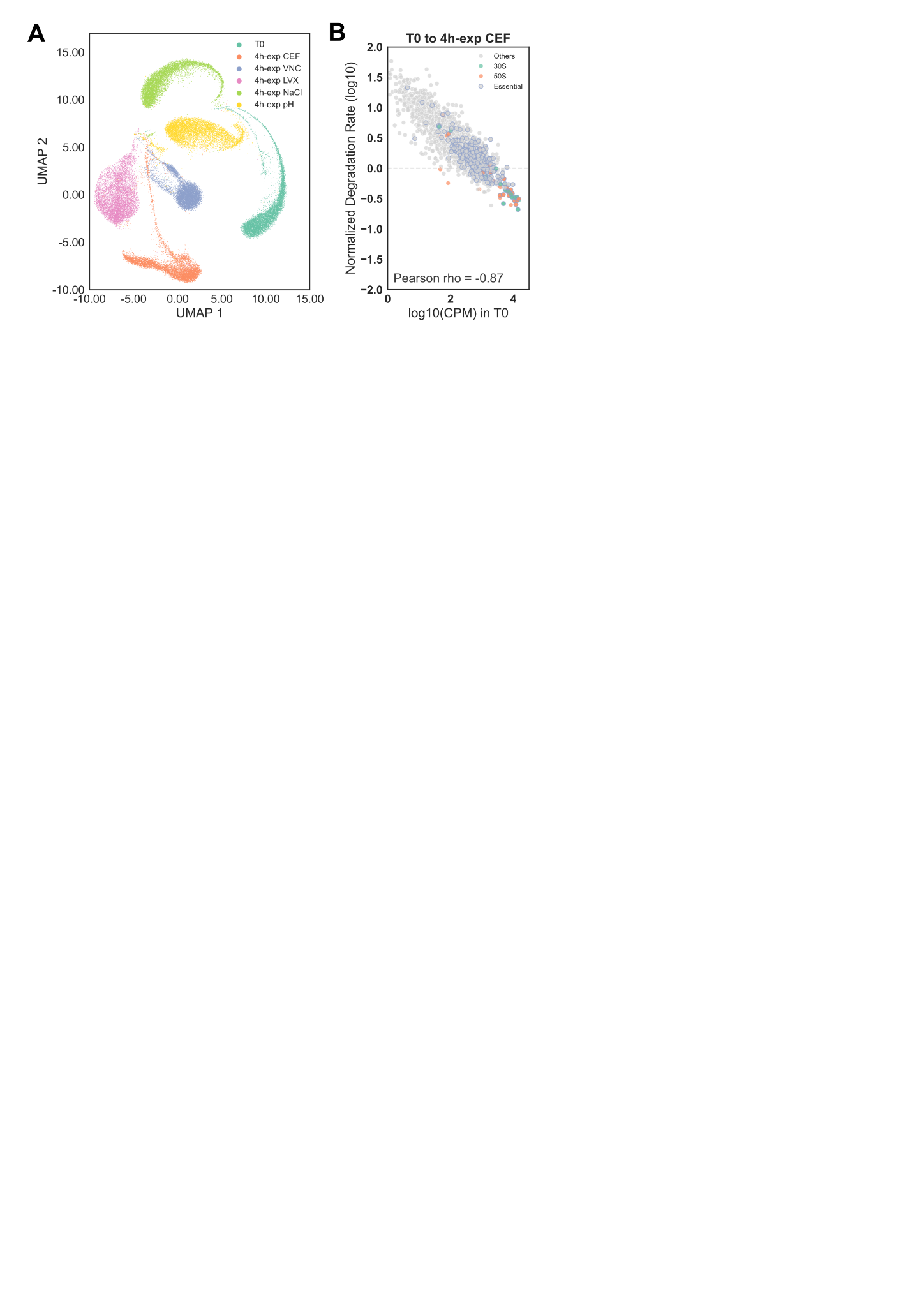
